# Masitinib mediates TGF-Beta1 and Nitric Oxide Secretion and Ameliorates MPTP/Microglia-Induced Degeneration of Differentiated SH-SY5Y Cells

**DOI:** 10.1101/2020.07.16.206094

**Authors:** Azize Yasemin Göksu Erol, Ersin Akıncı, Fatma Gonca Koçancı, Fatma Akçakale, Devrim Demir Dora, Hilmi Uysal

**Author notes:** Corresponding Author: Assoc. Prof. Dr. Azize Yasemin Göksu Erol, M.D., Dept. of Gene and Cell Therapy / Dept. of Histology and Embryology, School of Medicine, Akdeniz University, 07100, Antalya, Turkey.

## Abstract

**Introduction:** Microglia secretome includes not only growth factors and cytokines which support neuronal survival, it includes neurotoxic cytokines/enzymes, as well. MPTP is a neurotoxin which has degenerative effects on SH-SY5Y neuroblastoma cells. Masitinib mesylate is a tyrosine kinase inhibitor which has been shown to have beneficial effects in neurodegenerative diseases.

**Aim:** We first aimed to determine the most efficient microglial cell conditioned medium in terms of neurodegenerative effect. Next, we investigated the possible protective/therapeutic effects of masitinib against MPTP/microglia-induced degeneration of differentiated (*d*)-SH-SY5Y cells, and the role of transforming growth factor (TGF)-β1 and nitric oxide (NO) in these events.

**Material-Methods:** Non-stimulated/LPS-stimulated microglia cells were treated with masitinib or its solvent, DMSO. With or without MPTP-*d*-SH-SY5Y cell cultures were exposed to the conditioned media (CM) from microglia cell cultures, followed by cell survival analysis. Immunofluorescence staining of microglia and *d*-SH-SY5Y cells were performed with anti-CD-11b and anti-PGP9.5 antibody, respectively. TGF-β1/NO concentrations in CM of microglia/*d*-SH-SY5Y cell culture were measured.

**Results:** The initial 24 hrs CM of non-stimulated microglia cell culture was found to be the most detrimental microglial medium with lowest survival rates of treated *d*-SH-SY5Y cells. The toxicity of 48 and 72 hrs’ CM on *d*-SH-SY5Y cells were both lower than that of 24 hrs’ CM. Masitinib (0.5 µM), significantly prevented MPTP-related cell degeneration of *d*-SH-SY5Y cells. It also decreased the degenerative effects of both non-induced/LPS-induced microglia CM on with or without MPTP-*d*-SH-SY5Y cells. Although NO levels in microglia CM showed a negative correlation with survival rates of treated *d*-SH-SY5Y cells, a positive correlation was seen between TGF-β1 concentrations in microglial CM and rates of treated *d*-SH-SY5Y cell survival.

**Conclusion:** Masitinib ameliorates viability of with/without MPTP-*d*-SH-SY5Y cells. It does not only reverse the degenerative effects of its solvent, DMSO, but also prevents the degenerative effects of microglial secretions and MPTP. We suggest that masitinib begins to act as a neuroprotective agent via mediating TGF-β1 and NO secretion, as neurons are exposed to over-activated microglia or neurotoxins.

## 1. Introduction

Neurodegeneration, which can be defined as the progressive loss of functional neurons, is a common characteristic of Parkinson’s disease (PD), amyotrophic lateral sclerosis (ALS), Alzheimer’s disease (AD), and Huntington’s disease (HD). Although all of these are classified as neurodegenerative diseases, their underlying pathologies of central nervous system (CNS) are different. There is a large body of evidence for microglial activation in the pathogenesis of neurodegenerative disorders (1–3). Microglia are capable of secreting neuroprotective enzymes, such as superoxide dismutase 1 (SOD1) and lysozyme (4). On the other hand, negatively impacting neuronal viability, they secrete some neurotoxic enzymes such as matrix-metalloproteinases (MMPs), upon stimulation with lipopolysaccharide (LPS), a lipid-linked polymer of bacterial cell wall components found in Gram-negative bacteria (5–8). More importantly, LPS induced primary human microglia secrete some pro-inflammatory cytokines, such as interleukin 1 beta (IL-1β), IL-6, IL-8, Nitric oxide (NO), and tumor necrosis factor-alpha (TNF-α) (9).

Several cytokines, such as IL-10, IL-13, IL-1 receptor antagonist (IL-1RA), and transforming growth factor beta (TGF-β) are known to have anti-inflammatory activities by inhibiting excessive microglial activation in an autocrine and paracrine manner and also protect neurons from the toxic products secreted by microglia (10). NO is produced by macrophages in micromolar concentrations in response to inflammatory or mitogenic stimuli. Activated microglia, which are characterized with an amoeboid morphology, and increased expression of complement receptors and histocompatibility complex molecules, produce large amounts of cytotoxic molecules and pro-inflammatory cytokines (11,12).

Cell-cell interactions between microglia and neurons are mediated by e.g., cluster of differentiation 200 receptor (CD200R)-CD200 contact which provides constitutive inhibitory signals to microglia. Loss or disruption of these signals lead to a different microglial phenotype (13,14), which is characterized by the increased expression of activation markers, such as CD-11b and CD45 (15).

Multiple inhibitory signals from the CNS environment, including that of TGF-β, are important to maintain the microglial “M0” phenotype (14). Even under homeostatic conditions, microglia continuously sample the CNS environment with their highly motile processes and can hardly be described as “resting” but rather as neutral or “M0”. In this state, microglia are capable of protecting neurons from a variety of insults such as oxidative stress and excitotoxicity by releasing a subset of molecules. However, when microglial activation becomes chronic and unchecked as may be the case in several neurodegenerative disorders including AD and PD, these molecules may not be able to completely neutralize all neurotoxic secretions present in the CNS (5).

*In vitro* models are used to characterize microglial responses in responses against different pathologic stimuli and to test therapeutic interventions in such responses. *In vitro* cultures of neuronal cells are the basis for advancing knowledge about neurodegenerative diseases. In this context, *d-*SH-SY5Y neuroblastoma cell line is frequently used to study such diseases due to its neuronal properties with human origin, and ease of maintenance (16). 1-methyl-4-phenyl-1,2,3,6-tetrahydropyridine (MPTP) is a neurotoxin that triggers oxidative stress and apoptosis in neurons, making its usage possible in *in vivo* and *in vitro* neurodegenerative models (17).

Masitinib mesylate is a selective tyrosine kinase inhibitor that mainly targets type III growth factor receptors, including CD177 (tyrosine-protein kinase KIT; c-Kit), colony stimulating factor 1 receptor (CSF-1R), and platelet-derived growth factor receptors (PDGF-R) (18–20). It was demonstrated to be an efficient agent in controlling the survival, differentiation, and degranulation of mast cells (18–21). In addition to its therapeutic effects in cases of mastocytosis (21), the modulatory effect of masitinib on CNS function in multiple sclerosis (MS) (22), stroke (23), and AD (24) has also been shown. In this context, in a recent phase III clinical trial in which ALS patients were given masitinib, a reduction in the deterioration rate of motor functions has also been reported (25). Although the mechanisms of action related to masitinib remain largely unknown, the therapeutic effect of masitinib in ALS was attributed largely to the inhibition of CSF-1R in microglia and aberrant glial cells that surround degenerating motor neurons, and inhibition of mast cells via c-Kit receptors, as well (26–28). Masitinib is one of the most selective kinase inhibitors currently in development and it exerts a low toxicity profile (18). Trias et al. also reported that masitinib modulates the proliferation and inflammatory signaling underlying the emergence of aberrant glial cells, representing a new pharmacological approach to control detrimental neuroinflammation (20).

As *in vitro* models are instrumental to investigate cytokines and agents that may reverse degeneration and prevent loss of neurons, and since TGF-β is an important anti-inflammatory cytokine to maintain the microglial M0 phenotype, we aimed to evaluate the effects of LPS and masitinib on TGF-β1 production by microglia cells. We further examined the indirect/direct relation of masitinib to differentiated SH-SY5Y neuroblastoma cell viability, by means of microglial supernatant / direct application, respectively. We finally investigated the protective and/or therapeutic effects of masitinib on microglia-induced / MPTP related degeneration of human SH-SY5Y cells and the alterations of TGF-β1 levels in related conditions.

## 2. Material-Methods

### 2.1. Cell Culture and Group Characteristics

#### 2.1.1. HMC3 Microglia Cell Culture

The human embryonic microglia clone 3 cell line (HMC3) was purchased from ATCC (CRL-3304, Manassas, VA). (Note: This cell line was initially produced by transfecting human embryonic brain-derived macrophages with the large T antigen of the simian virus 40.).

HMC3 cells were cultured in Eagle’s Minimum Essential Media (EMEM) (11095-080-Thermo Fisher Scientific, Waltham, MA), supplemented with 10% fetal bovine serum (FBS) (04-007-1A-Biological Industries, Beit Haemenk, IL), non-essential aminoacids (11140-050-Gibco, Waltham, MA), and 1% antibiotic-antimycotic (L0010-020-Biowest, Nuaille, FR), and maintained in a humidified atmosphere, 5% CO_2_ at 37 °C.

#### 2.1.2. Group Characteristics of Microglia Cells

To determine the most potent neurodegenerative microglial cell culture supernatant/conditioned media (CM) and the effects of LPS (L2654-Sigma Aldrich, St. Louis, MO) and masitinib (13105-Cayman, Ann Arbor, MI) treatment on these cultures, HMC3 cells were grouped as four parts (Part 1-4), which were subsequently divided into subgroups [Table I].

Part 1 cells were stimulated with increasing doses of LPS (29).

Part 2-4 cells were treated with two different doses of masitinib; 0.5 µM low dose (M-LD) and 1 µM, high dose (M-HD) (20) in order to investigate the effects of masitinib on microglial activation and TGF-1β levels. Accordingly, Part 2-4 cells were treated with masitinib or its solvent, dimethyl sulfoxide (DMSO) (67-68-5-Merck EMSURE ACS) (1 µM), and supernatants were stored at -80 °C in order to be later applied on SH-SY5Y cells / be analyzed for TGF-β1/NO levels (Part 1’-4’).

Part 1-4 supernatants were later applied on **Part 1’-4’** d-SH-SY5Y cell groups [Table II].

##### *Treatment of <u>different doses of</u> <u>LPS</u> (Part 1) and* LPS-/+ and Masitinib/DMSO *(Part 2-4) on microglia cells*

Considering the importance of timing of the onset of the experiments after cell plating and the effects of duration of culture supernatant on its cytokine content, we followed 2 different protocols;

1. 24 hours (hrs) after plating the microglia cells, culture media were refreshed and then the experiment was initiated (Part 1 and 2).
2. The experiment was initiated as soon as the plated cells were attached (∼ 3 hrs) (Part 3 and 4) (Their supernatants were named below as ‘initial conditioned medium’)

Microglia cells were seeded in 6-well plates as 10^5^ cell/well. Groups were formed and the treatments described in Table I were performed.

After 24 hrs from the first treatment, all culture supernatants/CM were collected and stored at -80 ^°^C. To perform the morphological analysis and to measure the intensity of microglial activity, cells were stained with anti-CD11b antibody. Immunofluorescence (IF) analysis (Figure 6), morphological assessment and cell counting [Table III] were performed as described below [Table I-Part 1 and 2].

**Table I.** Group characteristics and treatment details of microglial cultures. *LPS and masitinib/DMSO were treated with an interval of 3 hrs in groups II- VI. ** All groups were cultured for 24 hrs.

| Part 1 | Part 2 | Part 3 | Part 4 |
| --- | --- | --- | --- |
| 24 hrs after plating microglia, cell media were refreshed, and groups were formed: |  | 3 hours after plating microglia (once cells attached), groups were formed. No media change. |  |
| I- No treatment<br>II- 1 µg/ml LPS<br>III- 5 µg/ml LPS<br>IV- 10 µg/ml LPS<br>V- 20 µg/ml LPS** | I- No treatment<br>II- M-HD followed by 10 µg/ml LPS<br>III- 10 µg/ml LPS followed by M-LD<br>IV- 10 µg/ml LPS followed by M-HD<br>V- DMSO followed by 10 µg/ml LPS<br>VI- 10 µg/ml LPS followed by DMSO*** | I- No treatment<br>II- M-LD<br>III- M-HD<br>IV- 1 µg/ml LPS<br>V- M-LD followed by 1 µg/ml LPS<br>VI- 1 µg/ml LPS followed by M-LD<br>VII- M-HD followed by 1 µg/ml LPS<br>VIII- 1 µg/ml LPS followed by M-HD<br>IX- DMSO followed by 1 µg/ml LPS<br>X-1 µg/ml LPS followed by DMSO** | I- No treatment- 24 hrs<br>II- 1 µg/ml LPS- 24 hrs<br>III- No treatment- 48 hrs<br>IV- 1 µg/ml LPS- 48 hrs<br>V- No treatment- 72 hrs<br>VI- 1 µg/ml LPS- 72 hrs |
| Microglia were stained with anti-CD-11b antibody, IF analysis, cell count, morphological assessment were performed. |  |  |  |
| After 24 hours from the first treatment, CM of all groups were collected. |  |  | After 24/48/72 hrs from the first treatment CM of all groups were collected. |

**Figure 1.**
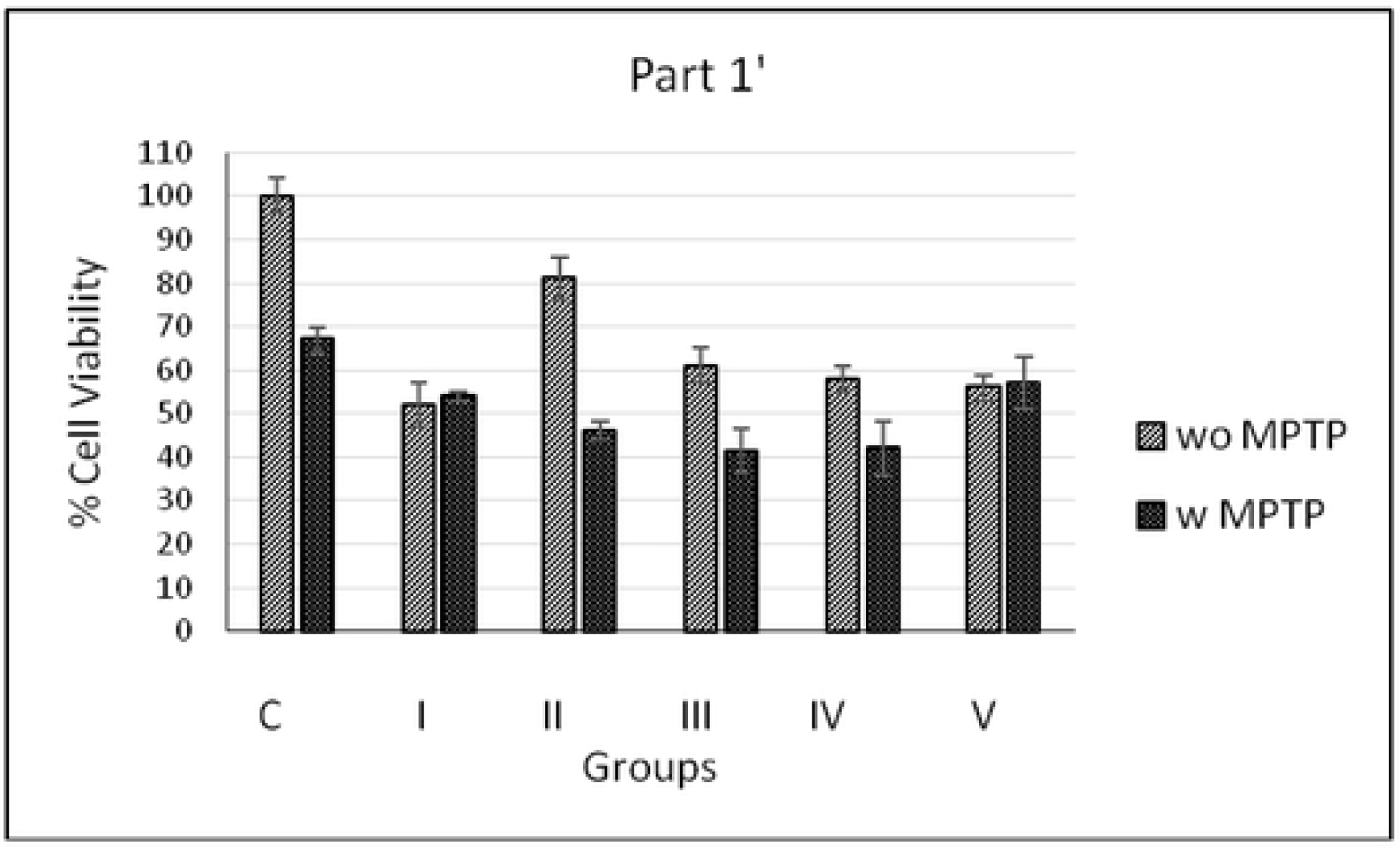
Effects of CM from microglia culture stimulated with different doses of LPS on viability of *d-*SH- SY5Y cells with or without MPTP. **C:** Control (Non-CM treated *d-*SH-SY5Y cell) **I:** No LPS, **II:** 1 µg/ml LPS, **III:** 5 µg/ml LPS, **IV:** 10 µg/ml LPS, **V:** 20 µg/ml LPS -induced microglia CM treated *d-*SH-SY5Y cell As seen in the Figure 1, MPTP treatment decreased the cell viability at a rate of about 35% (C)*. *Included in all graphics.

#### 2.1.3. Human SH-SY5Y Neuroblastoma Cell Culture and Differentiation

SH-SY5Y cells were obtained from SAP Institute (Ankara, Turkey). The cells were maintained in Dulbecco’s Modified Eagle’s Medium (DMEM) (D5796-Sigma Aldrich) supplemented with 10% FBS (10500064-Gibco), 1% penicilin/streptomycin (15140122-Sigma Aldrich), 1% L-glutamine (25030024-Sigma Aldrich), 40% MCDB-201 (M6770-Sigma Aldrich), and cultured in a humidified CO_2_ (5%) incubator at 37 °C. The medium was changed every other day. For neuronal differentiation, SH-SY5Y cells were treated with 10 µM Retinoic acid (RA) (R2625-Sigma) once every other day in DMEM with 2% FBS for 6 days. The differentiated-SH-SY5Y (*d-*SH-SY5Y) cells with acquired neuronal properties were further used in experiments.

1-methyl-4-phenyl-1,2,3,6-tetrahydropyridine (MPTP) (M0896-Sigma Aldrich), a neurotoxin, was used to degenerate *d-*SH-SY5Y cells. To find the most appropriate MPTP dose which degenerates 30-40% of *d-*SH-SY5Y cells, we performed dose titration assay for cell viability, including 5 different doses up to 1500 µM. As 1000 µM MPTP was found to be the most appropriate dose for degeneration causing around 35% of cell death, this dose of MPTP was further used in all our experiments to degenerate *d-*SH-SY5Y cells.

#### 2.1.4. Group Characteristics of *d-*SH-SY5Y Cells

In order to determine the degenerative effects of microglial CM, with or without MPTP, *d-*SH-SY5Y cells in Part 1’-4’ were treated with these CM, followed by 3-[4,5-dimethylthiazol-2-yl]-2,5-diphenyltetrazolium (MTT) (ab146345-Abcam, Cambridge, UK) assay. On the other hand, to determine the protective/therapeutic effects of masitinib against MPTP-related cell degeneration, *d-*SH-SY5Y cells were treated with masitinib which is followed by MTT assay and IF analysis in Part 5’ and 6’, respectively. *Note: Masitinib solvent, DMSO was also applied*.

#### Groups as *Part 1’-6’* were formed and treatments were performed as described in Table II

##### *Treatment of* Microglia CM Part 1-4 *and* Masitinib/DMSO *on with or without MPTP-d-SH-SY5Y cells*

**Part 1’-4’**: *d-*SH-SY5Y cells were plated in 96-well plates (5×10^3^ cell/well), cultured for 48 hrs, followed by MPTP treatment to half of the plates. with or without MPTP-*d*-SH-SY5Y cells were exposed to Part 1-4 supernatants 1 hour after MPTP treatment to investigate the effects of different groups of microglia CM on these cells. After 24 hrs from the last treatment, cell survival analysis was performed with MTT assay (Figures 1-4).

**Part 5’**: *d*-SH-SY5Y cells were plated in 96-well plates (5×10^3^ cell/well), cultured for 48 hrs, followed by MPTP treatment of half of the plates. Part 5’ with 5 subgroups were formed. with or without MPTP-*d*-SH-SY5Y cells were exposed to Masitinib/DMSO before (early) or after (late) MPTP treatment to investigate the protective and therapeutic effects of masitinib.

*** Early: 24 hrs before MPTP treatment, Late: 1 hr after MPTP treatment [Table II]. After 24 hrs from the last treatment, cell survival analysis was performed with MTT assay (Figure 5).

**Part 6’**: *d*-SH-SY5Y cells were seeded as 25⨯10^3^ cell/well in 24-well plates. Part 6’ with 11 subgroups were formed [Table II]. with or without MPTP-*d*-SH-SY5Y cells were exposed to various microglia CM (with or without masitinib) or treated with M-HD. 24 hrs later, IF staining with anti-PGP9.5 antibody was performed, and fluorescence microscopy images of *d-*SH-SY5Y cells were taken (Figure 8). Morphological analysis (Figure 7), and cell degeneration rates were scored [Table IV].

**** M-HD/CM treatments were performed 1 hr after MPTP treatment [Table II].

**Table II.** Group characteristics and treatment details of with or without MPTP- *d-*SH-SY5Y cell cultures.

| Part 1' | Part 2' | Part 3' | Part 4' | Part 5' | Part 6' |
| --- | --- | --- | --- | --- | --- |
| *with or without MPTP- d-SH-SY5Y cells (5x10<sup>3</sup>/well) in 96-well plates* |  |  |  |  | *d-SH-SY5Y cells (25x10<sup>3</sup>/well) in 24-well plates* |
| Treated with Part 1 microglia CM | Treated with Part 2 microglia CM | Treated with Part 3 microglia CM | Treated with Part 4 microglia CM | I- Early M-LD<br>II- Late M-LD<br>III- Early M-HD<br>IV- Late M-HD<br>V- Early DMSO *** | I- MPTP (-) (Control)<br>II- MPTP (+)<br>III- MPTP (+), Masitinib (1 μM)<br>IV- 24 hrs non-induced microglia CM<br>V- MPTP (+), 24 hrs non-induced microglia CM<br>VI- MPTP (-), 24 hrs 1 μg/ml LPS-induced microglia CM<br>VII- MPTP (+), 24 hrs 1 μg/ml LPS-induced microglia CM,<br>VIII- MPTP (-), 24 hrs 10 μg/ml LPS-induced microglia CM<br>IX- MPTP (+), 24 hrs 10 μg/ml LPS-induced microglia CM,<br>X- MPTP (-), 24 hrs 10 μg/ml LPS-induced, masitinib treated microglia CM,<br>XI- MPTP (+) 24 hrs 10 μg/ml LPS-induced, masitinib treated microglia CM**** |
| MTT assay was performed. |  |  |  |  | IF, morphological assessment, and cell counting were performed. |

### 2.2. Methods

#### 2.2.1. Cell Viability Assay

Cell viability analysis was performed with MTT assay in Part 1’-5’ *d*-SH-SY5Y cell culture groups [Table II, Figs 1-5]. *d-*SH-SY5Y cells were plated into 96-well microplate (5⨯10^3^ cell/well). After 24 hrs incubation at 37°C with 5% CO_2_, MPTP and/or other drug/chemical and supernatant treatments were performed. After an additional 24 hrs of culture, MTT, membrane-permeable dye solution was added to the cells with a final concentration of 1 mg/mL, and the mixture was incubated at 37 °C for about 4 hrs. The supernatant was removed, then the pellets were dissolved in 200 μl/well of DMSO. The absorbance was measured by spectrophotometry at 570 nm using microplate reader (Thermo Scientific Multiskan Spectrum). Viability was accepted as the percentage of absorbance of treated cells to non-treated control cells (30).

#### 2.2.2. Immunofluorescence (IF) Assay

HMC3 and SH-SY5Y / *d-*SH-SY5Y cells were immunostained with anti-CD-11b (nb11089474 pAB-Novus, Boston, MA) and anti-PGP9.5 (ab1503-Abcam) antibody, respectively.

The cells were fixed with paraformaldehyde for 20 min. and blocked with phosphate buffered saline (PBS) containing 0.1% Tween-20 (v/v) and 5% bovine serum albumine for 1 hr at room temperature.

The HMC3 cells were immunostained with anti-CD-11b antibody (1:400) at 4 °C overnight, and then incubated with secondary antibody (35552-Goat Anti-Rabbit IgG, DyLight 488, Thermo Scientific) (1:400) and DAPI (ab104139-Abcam).

SH-SY5Y / *d-*SH-SY5Y cells were immunostained with anti-PGP9.5 antibody (1:600) at 4 °C overnight, and then incubated with secondary antibody (A10037-Alexa Fluor 568 donkey anti-mouse IgG (H+L)- Invitrogen) (1:500) and DAPI.

Fluorescent intensity was evaluated and photographs were taken using a Leica DMi8 Microscope and LasX software (Figures 6-8).

#### 2.2.3. Cell Counting and Morphological Analysis

##### 2.2.3.1. Assessment of Microglia

Microscopic evaluation of microglia cell groups included these parameters: Cell counting of total/amoeboid cells and intensity of staining with anti-CD-11b antibody [Fig 6; Table III].

Microglia cells were counted by staining these cells with antibody against CD11b, which is a marker commonly used to evaluate microglial activation. Cytoplasmic staining intensity was scored 0 to 5. Microglia cell counts (MCCs) and amoeboid cell counts (ACCs) were determined as follows.

First, the area with highest microglial cell number was identified at low power (40× and 100×). MCCs were then performed on a 200× field. The mean number of MCCs was calculated from images of 5 different area. On the other hand, the mean number of ACCs per 5 fields with the most abundant infiltration at a magnification of ×400 was calculated for each group.

##### 2.2.3.2. Assessment of d-SH-SY5Y cells

Microscopic evaluation of *d-*SH-SY5Y cell groups included these parameters: Overall approximate cell count, numbers of large, medium, and scattered small cell aggregates, the rate of degeneration inside large aggregates (scored from 0 to 10 according to the signs of cell degeneration/necrosis, including recognizable structural changes, abnormal accumulations in cytoplasm with granular appearance, fragmented / pyknotic nuclei, diminished cell size, and lysis of cell membrane) [Figs 7, 8; Table IV].

#### 2.2.4. TGF-β1 (Enzyme-Linked Immunosorbent Assay (ELISA) Assay and Nitrate/Nitrite Colorimetric Assay

Cell-free culture supernatants of microglia and *d-*SH-SY5Y cells were collected and assayed for cytokine TGF-β1 with ELISA using Human/Mouse TGF-β1 Elisa kit (88-8350-eBioscience). NO levels in some microglia CM were measured with Nitrate/Nitrite Colorimetric Assay Kit (78001-Cayman, MI, USA) according to the manufacturer’s instructions. Nitrate and nitrite present in culture supernatants (80 μl) were converted to NO.

Note: Very high NO levels lead to the formation of peroxynitrite and destruction of iron-sulfur clusters. The final products of NO in vivo are nitrite and nitrate.

## 3. Results

### 3.1. Cell Viability

#### 3.1.1. The Effects of Non-Stimulated or Increasing Doses of LPS-Stimulated Microglia CM on *d-*SH-SY5Y Cell Viability

These results belong to the 1/4 ratio of microglia CM / *d-*SH-SY5Y cell culture medium. As we applied different concentrations (1/2, 1/4, 1/8) of microglia CM on *d-*SH-SY5Y cells, we found that cell viability rates show a negative correlation with increasing concentrations of the CM applied (Data not shown).

In without MPTP- *d-*SH-SY5Y cells, microglia CM which were not activated with LPS (Group P1’- I) caused a higher rate of cell degeneration when compared to the ones activated by increasing doses of LPS (Groups P1- II-V). When the latter supernatants were applied, the dose of LPS that microglia were exposed showed a negative correlation with the viability of SH-SY5Y cells (Groups P1’- II-V).

In MPTP-treated *d-*SH-SY5Y cells, degeneration rate was lower in cells that exposed to non-induced microglia supernatants (Group P1’- I) compared to 1, 5, 10 µg/ml LPS-treated microglia supernatants (Groups P1’- II-IV). However 20 µg/ml LPS-treated microglia supernatant (Group P1’- V) caused a similiar degeneration with that of non-treated microglia supernatant (Group P1’- I) (Figure 1).

#### 3.1.2. The Effects of 10 µg/ml LPS-Stimulated, Masitinib-Treated Microglia CM on *d-*SH-SY5Y Cell Viability

The treatment of *d-*SH-SY5Y cells with microglia CM in which masitinib administration was performed before (Group P2’- II) or after (Low dose: Group P2’- III, High dose: Group P2’- IV) LPS stimulation resulted in a lesser percentage of cell death when compared to non-masitinib treated group (Group P2’- I). Microglia CM that exposed to Masitinib-solvent, DMSO significantly decreased cell viability. There was no significant difference between the effects of DMSO treatment before or after LPS on cell viability (Groups P2’- V, VI, respectively) (Figure 2).

**Figure 2.**
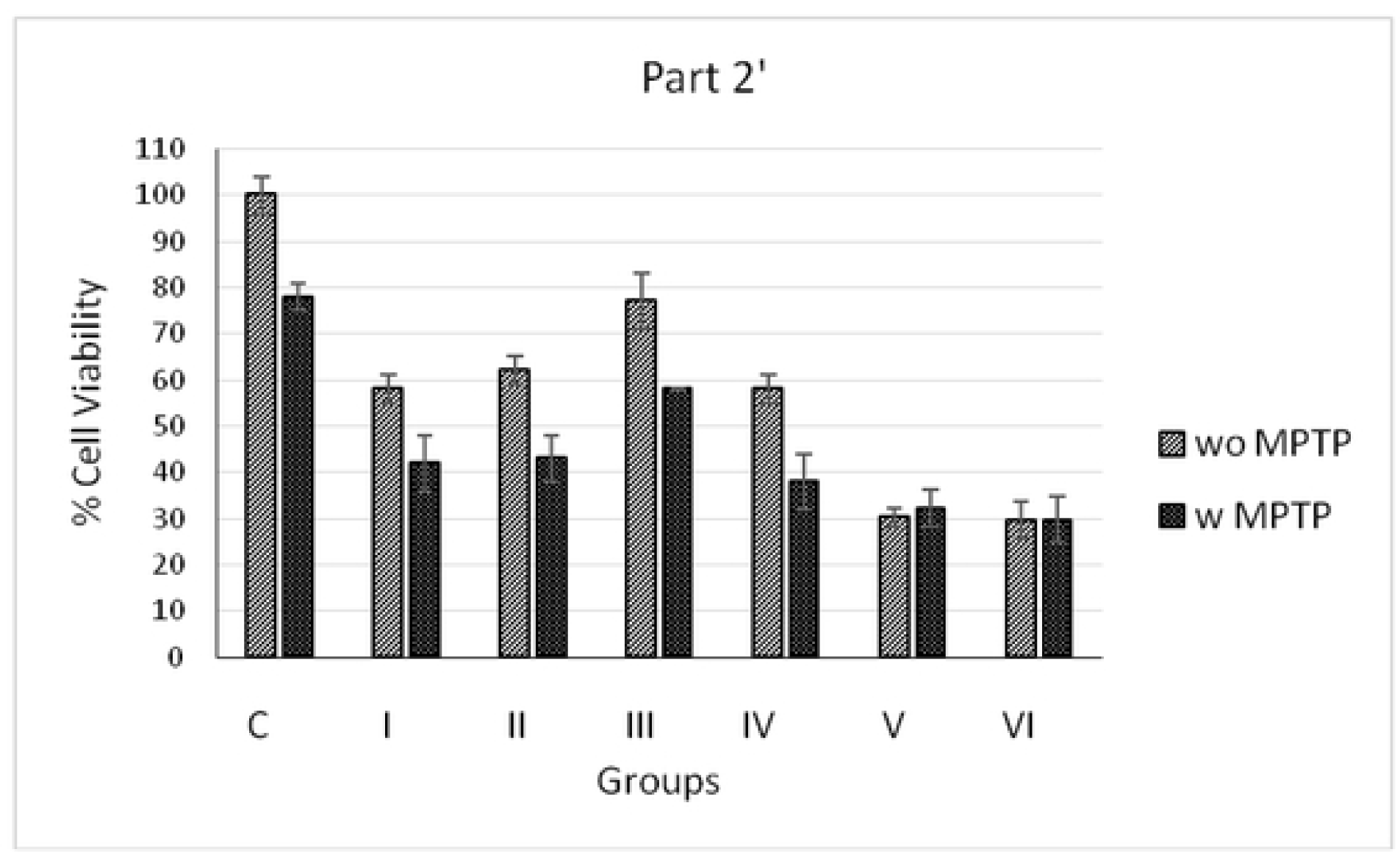
Effects of microglia CM in which masitinib/DMSO administration were performed before or after 10 µg/ml LPS stimulation on *d-*SH-SY5Y cell viability. C: Control (Non-CM treated *d-*SH-SY5Y cell) **I:** No LPS, **II:** M-HD + 10 µg/ml LPS, **III:** 10 µg/ml LPS + M-LD, **IV:** 10 µg/ml LPS + M-HD, **V:** DMSO + 10 µg/ml LPS, **VI:** 10 µg/ml LPS + DMSO -administered microglia CM were treated on *d-*SH-SY5Y cells

#### 3.1.3. The Effects of 1 µg/ml LPS-Stimulated, Masitinib-Treated Initial Microglia CM on *d-*SH-SY5Y Cell Viability

The initial 24 hrs’ CM of microglia which were not activated with LPS (Group P3’- I) caused significant degeneration (only around 5% cell viability) on both with and without MPTP *d-*SH-SY5Y cells when compared to control group. This initial non-LPS-stimulated microglia CM was obviously the most detrimental medium for both with or without MPTP *d-*SH-SY5Y cells. Importantly, M-LD and M-HD treated microglia CM (Groups P3’- II and III, respectively) showed less degenerative effects on d-SHSY5Y cells when compared to non-treated ones (Group P3’- I).

The initial 24 hrs’ CM of microglia which were activated with 1 µg/ml LPS caused also significant degeneration, with around 26% cell viability of both with and without MPTP *d-*SH-SY5Y cells exposed to this CM (Group P3’- IV). This initial 1 µg/ml-LPS-stimulated microglia CM was the second most detrimental medium –following initial medium of non-stimulated microglia- for both with and without MPTP d-SH-SY5Y cells. Importantly, both early and late administration of M-LD (Group P3’- V, VI, respectively) and M-HD (Group P3’- VII, VIII, respectively) ameliorated the degenerative effects of this supernatant. Late M-HD (Group P3’- VIII) showed lower cell viability rates, compared to that of Early M-HD (Group P3’- VII), and Early and Late M-LD (Group P3’- V-VI), in both with and without MPTP *d-*SH-SY5Y cells.

The treatment of *d-*SH-SY5Y cells with microglial supernatants in which masitinib-solvent, DMSO administration was performed on stimulated / non-stimulated microglia (Groups P3’- IX and X, respectively) resulted in a lesser percentage of cell viability when compared to masitinib treated ones (Figure 3).

**Figure 3:**
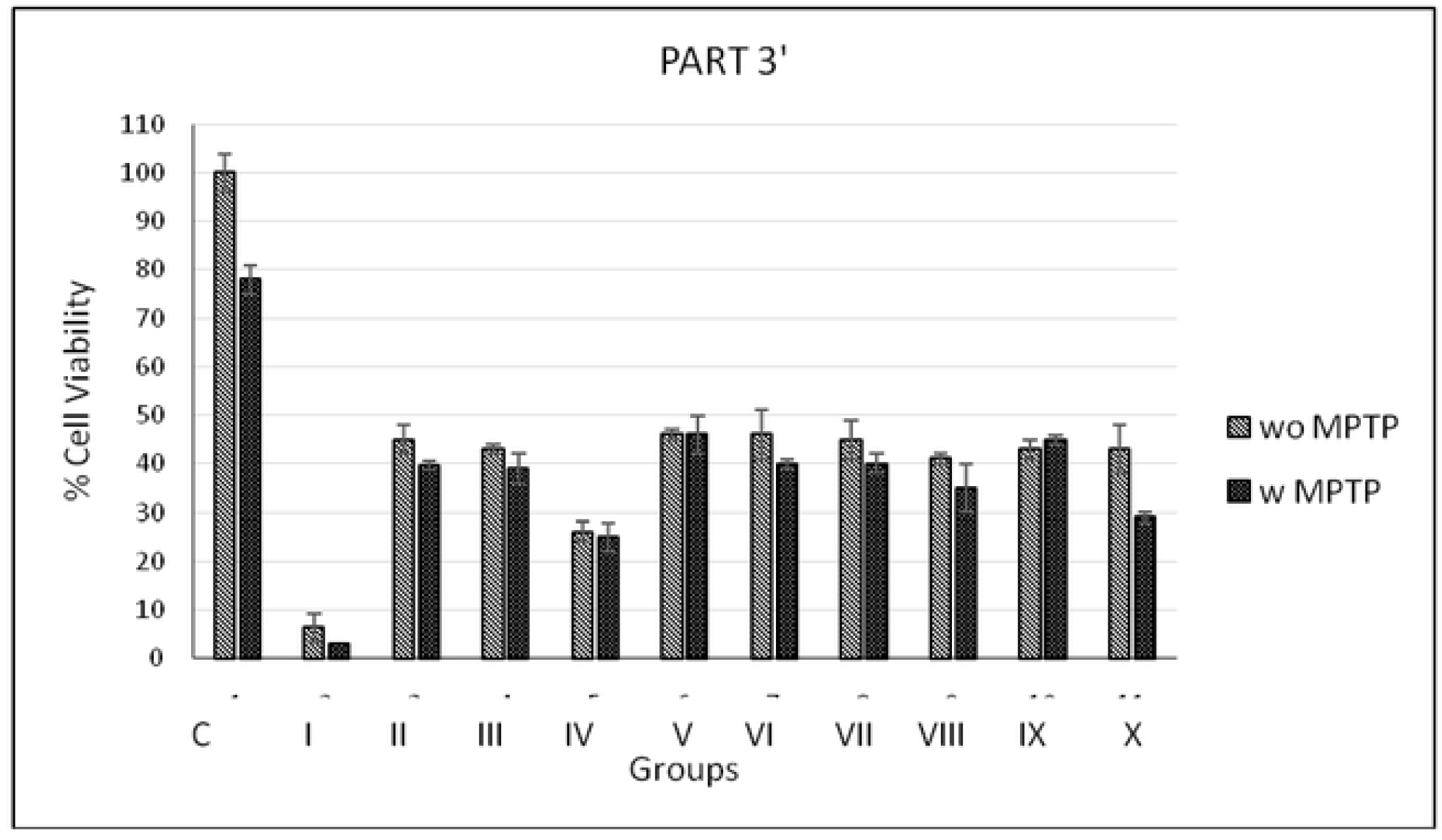
Effects of initial 24 hrs microglia CM -in which masitinib/DMSO administration were performed before or after 1 µg/ml LPS stimulation-on *d-*SH-SY5Y cell viability. **C**: Control (Non-CM treated *d-*SH-SY5Y cell) **I:** Initial 24 hrs CM (not activated with LPS), **II:** M-LD, **III:** M-HD, **IV:** 1 µg/ml LPS, **V:** M-LD + 1 µg/ml LPS, **VI:** 1 µg/ml LPS + M-LD, **VII:** M-HD + 1 µg/ml LPS, **VIII:** 1 µg/ml LPS + M-LD, **IX:** DMSO + 1 µg/ml LPS, **X:** 1 µg/ml LPS + DMSO -administered microglia CM which were treated on *d-*SH-SY5Y cells)

#### 3.1.4. The Effects of 24, 48, and 72 hours’ Culture of Non-stimulated or 1 µg/ml LPS-Stimulated Microglia CMs on *d-*SH-SY5Y Cell Viability

The initial 24 hr CM at high concentration (1/2 ratio) of microglia which were not activated with LPS caused very significant degeneration, with around 6% cell viability of *d-*SH-SY5Y cells (Group P4’- I). 48 hrs CM of non- stimulated microglia (Group P4’- III) were significantly less detrimental on *d-*SH-SY5Y cells with better viability rates, compared to that of 24 hrs Group P4’-I. However, 72 hrs CM treatment caused a moderate effect.

The initial CM of 1 µg/ml-LPS-stimulated microglia at 24, 48 and 72 hr (Groups P4’- II, IV and VI, respectively) caused better viability rates on with or without MPTP- d-SH-SY5Y cells, when compared to non LPS-stimulated ones, and the viability rates of *d-*SH-SY5Y cells were found to be increased when exposed to CM of prolonged culture (Figure 4).

**Figure 4.**
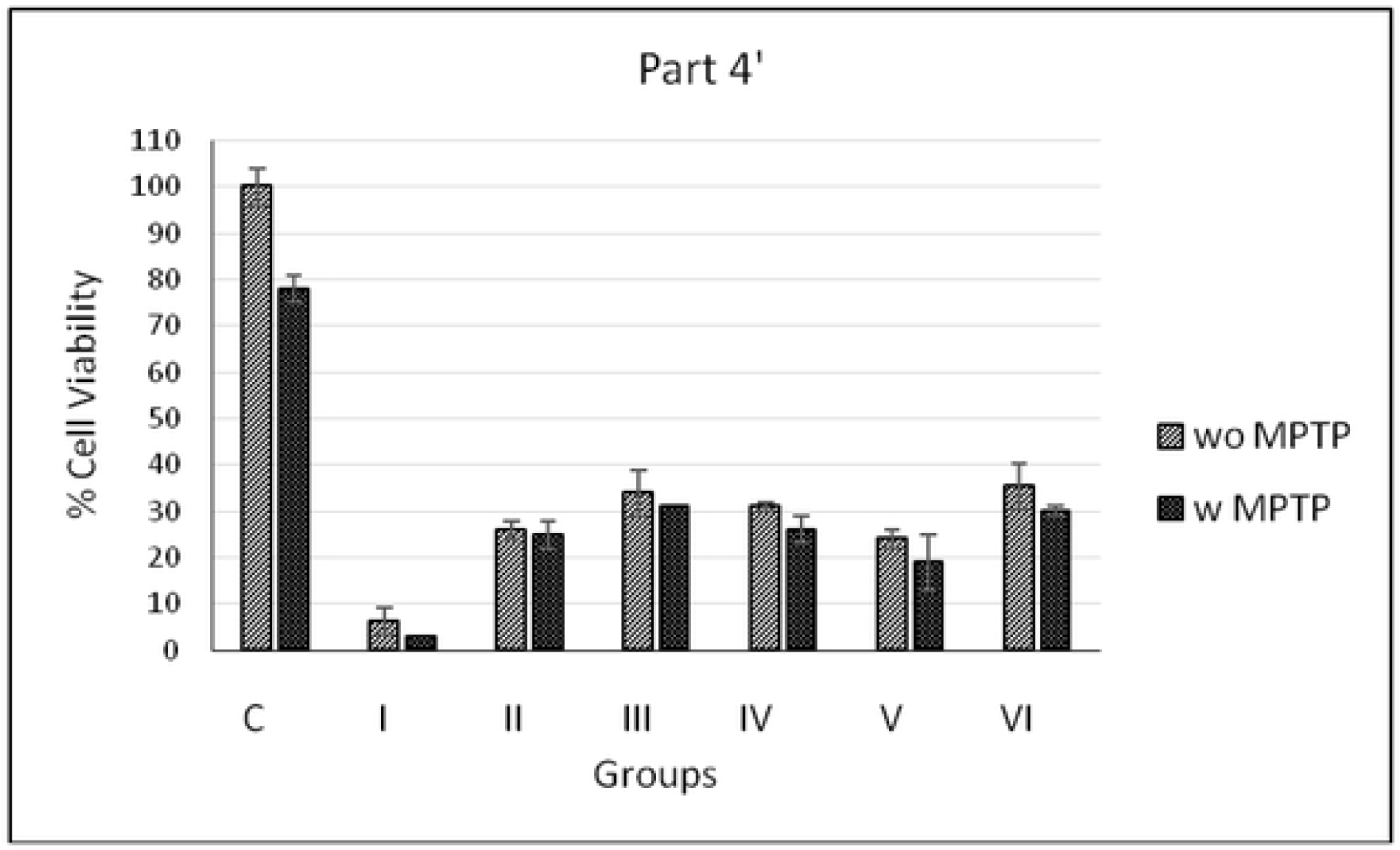
Effects of 24, 48, and 72 hrs’ microglia CM (n/a or activated with LPS) on *d-*SH-SY5Y cell viability C: Control (No CM treatment *d-*SH-SY5Y cell) **I:** No LPS-24 hrs, **II:** 1 µg/ml LPS-24 hrs, **III:** No LPS-48 hrs, **IV:** 1 µg/ml LPS-48 hrs, **V:** No LPS-72 hrs, **VI:** 1 µg/ml LPS-72 hrs -cultured microglia CM treatment on *d-*SH-SY5Y cells

#### 3.1.5. The Effect of pre- or post- Masitinib Treatment on Viability of *d-*SH-SY5Y Cells Exposed to MPTP, Hydrogen Peroxide or Microglial CM

When M-HD (Group P5’- III), and DMSO 1 µM (Grup P5’- V) were administered 24 hrs before MPTP treatment, cell viability rates were found to be lower than control, but M-LD (Grup P5’- I) did not cause such a decrease in viability rates. Importantly, late administration of M-LD (Group P5’- II) significantly ameliorated the viability rates of MPTP treated *d-*SH-SY5Y cells, reaching their viability rate to 100% (Figure 5).

**Figure 5.**
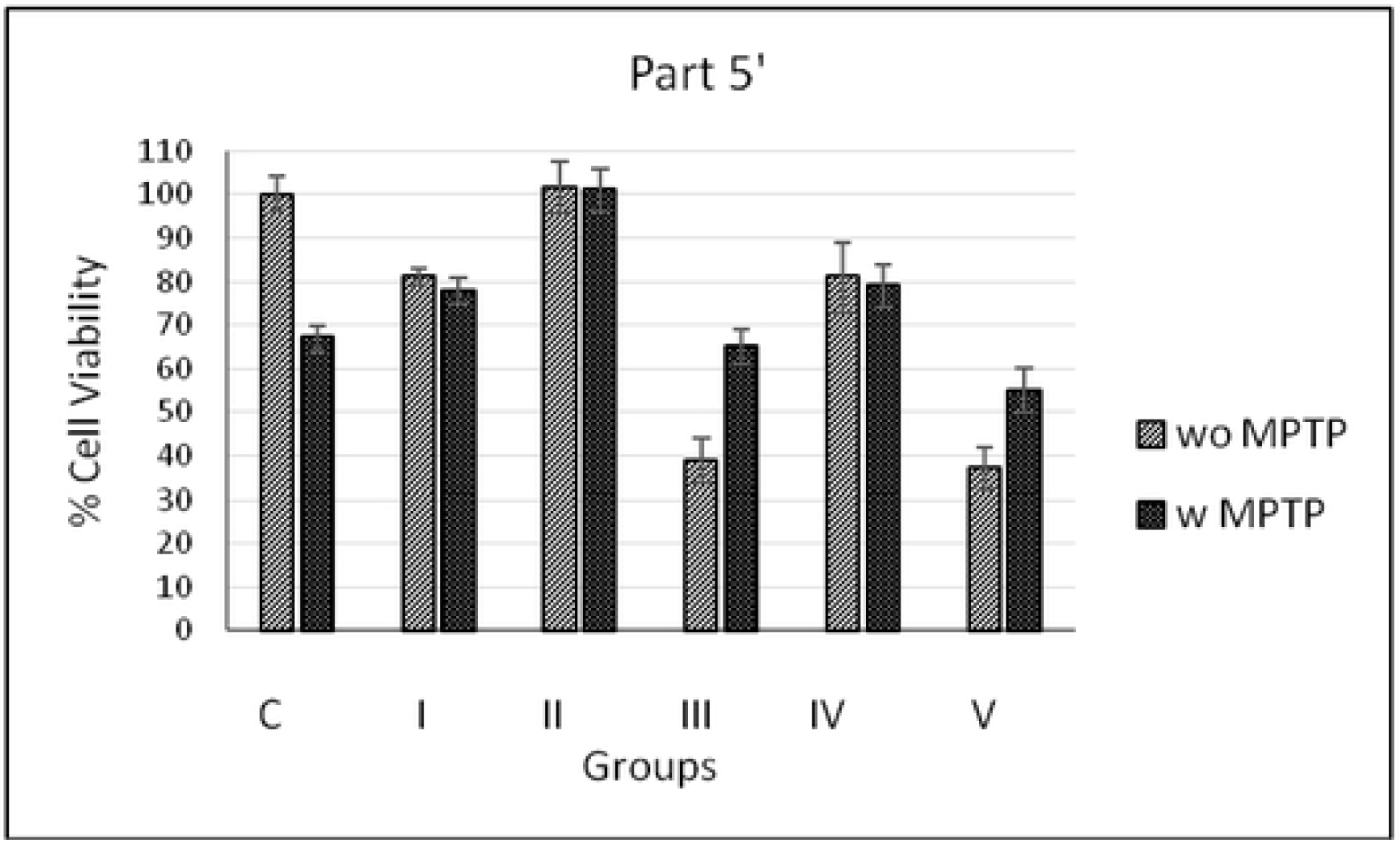
Effects of masitinib (early/late) treatment on with or without MPTP *d-*SH-SY5Y cell viability C: Control (Non-treated *d-*SH-SY5Y cell) **I:** Early M-LD **II:** Late M-LD, **III:** Early M-HD, **IV:** Late M-HD, **V:** Early DMSO -treated with or without MPTP *d-*SH-SY5Y cells.

In addition to MPTP, we also investigated the protective effects of M-LD against degeneration caused by hydrogen peroxide and initial 24 hrs’ microglial CM, as well. In both degenerative conditions, M-LD showed its protective effect by increasing survival rates of treated *d-*SH-SY5Y cells, with a similiar rate of protection that it showed against MPTP.

### 3.2. IF analysis

#### 3.2.1. IF Images of Microglia Cells

In control group with no LPS, the average number of microglia cells, a few number of ameboid microglia cells, and medium intensity of Cd-11b staining were observed. In 1 ug/ml LPS treated group, while microglia cell number and intensity of Cd-11b staining remained similar, the number of ameboid microglia cells modestly increased. Starting from 5 ug/ml LPS treated group up to 20 ug/ml LPS group, all three parameters demonstrated a significant and consistent increase; high number of microglia and ameboid microglia cells as well as high Cd-11b expression. On the other hand, 10 ug/ml LPS and masitinib treated groups demonstrated a decrease both in the number of microglia cells, and ameboid microglia cells as well as intensity of Cd-11b staining. Whereas, LPS/DMSO and DMSO/LPS treated groups did not show any improvement compared to their masitinib treated counterparts (Figure 6). The data regarding to changes in cell number and intensity of IF stained Cd-11b proteins upon LPS and/or masitinib treatment were also summarized in Table III.

**Figure 6.**
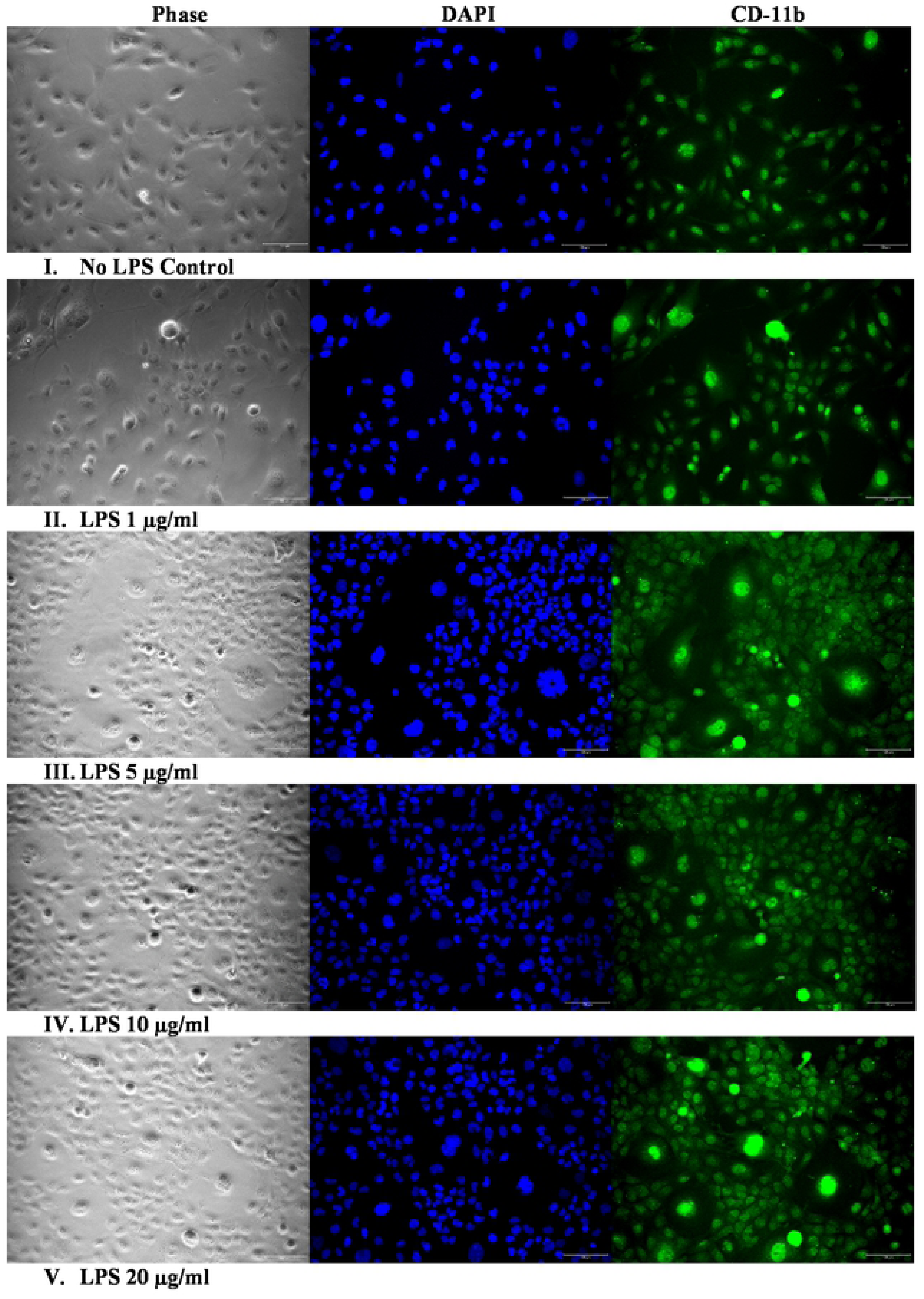
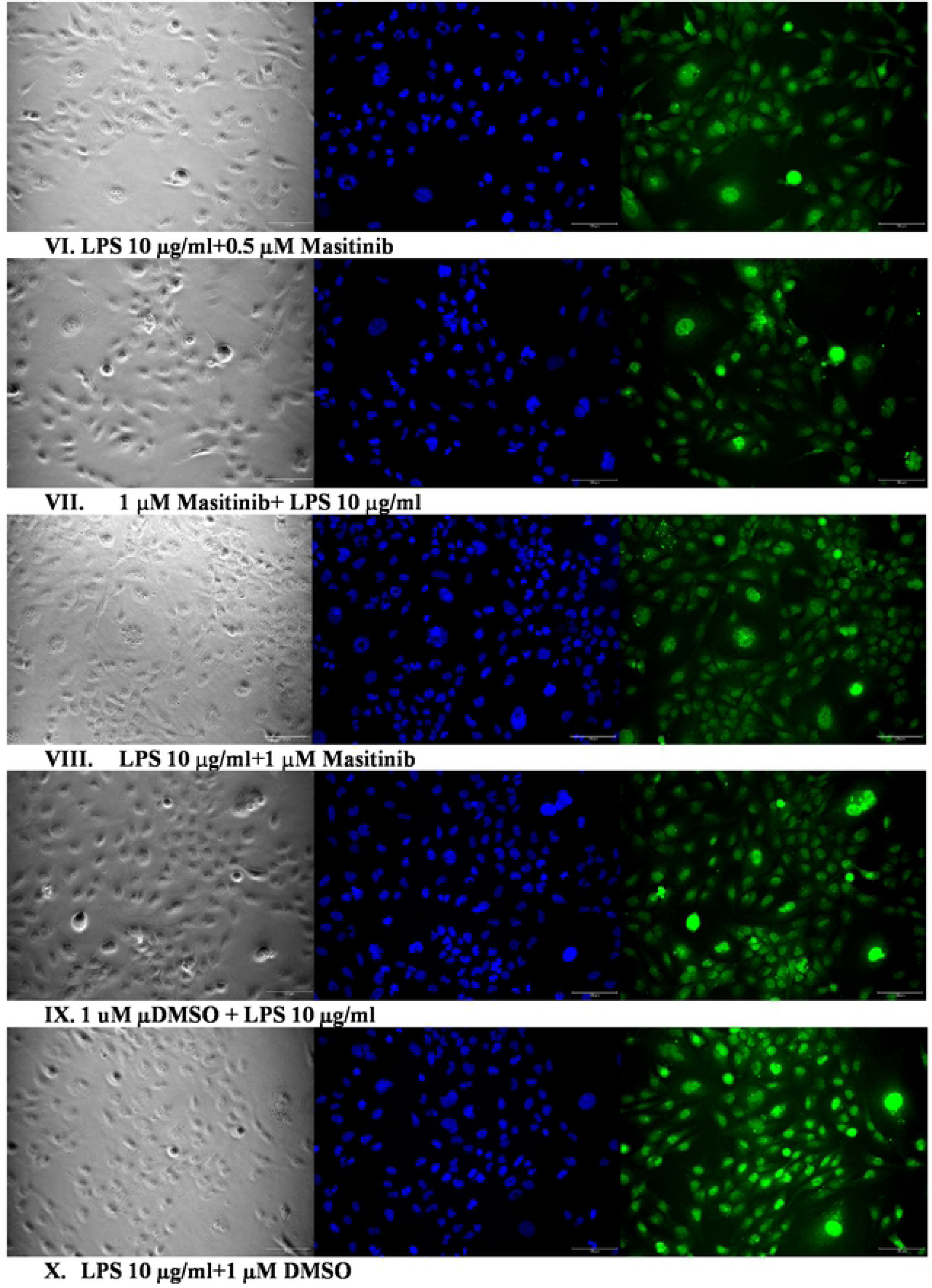
IF microscopy images of microglia cells treated with differant concentration of LPS and/or Masitinib. DAPI is blue, CD-11b is GFP. Scale bar is 10 uM.

#### 3.2.2. IF Images of SH-SY5Y Cells

In MPTP-treated *d*-SH-SY5Y cells (II), PGP9.5 staining intensity was found to be lower than that of control (I). In MPTP + M-HD group (III), PGP9.5 staining intensity was significantly increased compared to group II. Moreover, when compared to treatment with LPS-induced microglia CM (IV), M-HD + LPS-induced microglia CM caused higher intensity of PGP9.5 staining in *d*-SH-SY5Y cells (Figure 7).

Immunostaining of *d-*SH-SY5Y cells with anti-choline acetyltransferase (ChAT) antibody, a cholinergic marker was also performed, in which *d-*SH-SY5Y cells showed an increased ChAT activity in response to RA treatment.

**Figure 7.**
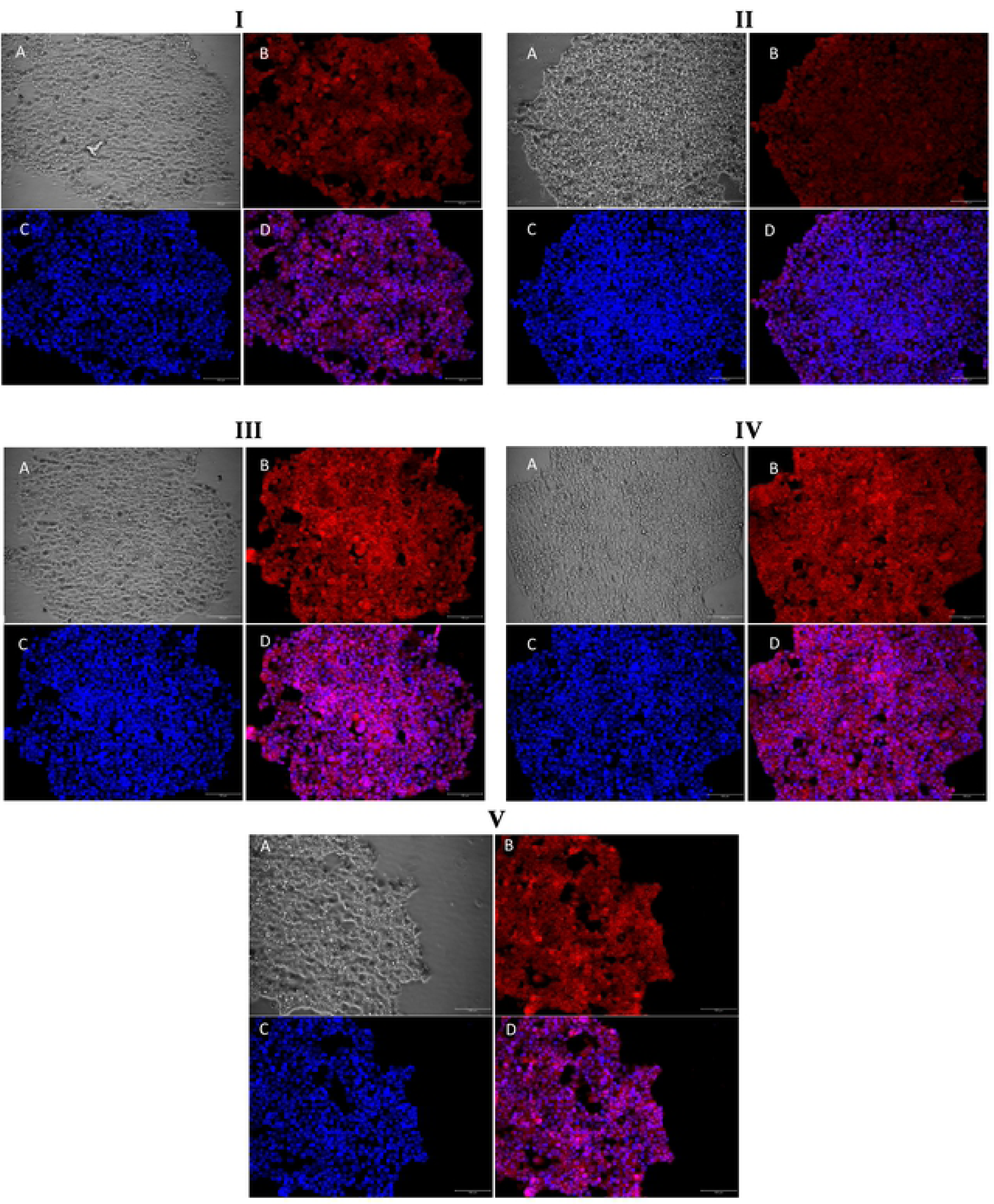
Fluorescence microscopy images of immunostained SH-SY5Y cells. **A:** Phase, **B:** PGP9.5 (red), **C:** DAPI (blue), **D:** Merge (red-blue) (x200) **I:** Non treated, **II:** MPTP-treated, **III:** MPTP + Masitinib High Dose-treated, **IV:** 10 µg/µl LPS-induced microglia supernatant treated, **V:** Masitinib High Dose-applied 10 µg/µl LPS-induced microglia supernatant treated.

### 3.3. Cell count and Morphological analysis

#### 3.3.1. Microglia cells

In microglia cell cultures that were exposed to 1, 5 and 10 µg/ml LPS (II-IV), microglial cell numbers were found to be increased in parallel to LPS dose. However, 20 µg/ml LPS (V) did not cause such an increase in microglial number. In the latter group, amoeboid cell number was found to be significantly increased compared to all other groups. Thus, these results may explain the possible cause of higher rates of cell viability that was seen in *d-*SH-SY5Y cells exposed to CM of 20 µg/ml LPS induced microglia (V).

In ‘LPS + M-LD treated microglia group (VI), microglial cell number and CD-11b intensity were found to be lower than that of M-HD treated one (VIII). This may explain the higher rates of cell viability seen in *d-*SH-SY5Y cells that were exposed to M-LD treated microglia CM compared to M-HD treated one.

**Table III.**
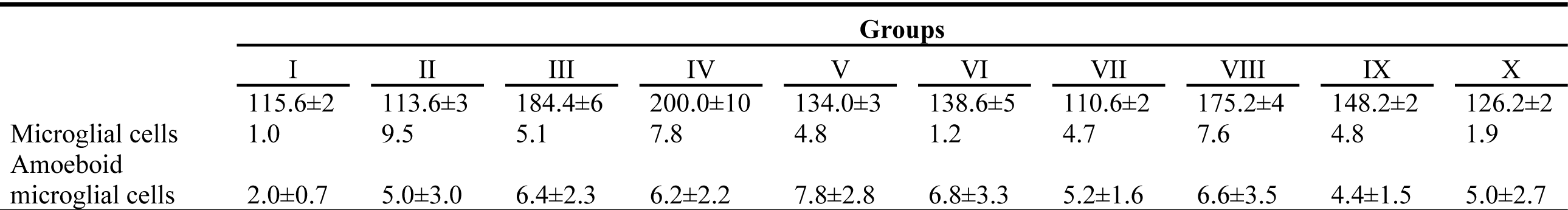
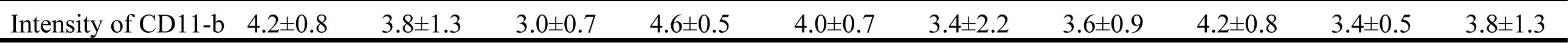
Cell count of microglia and amoeboid cells, and staining intensity with CD11b antibody (24 hrs’ culture). **I-**No treatment, **II-** 1 µg/ml LPS, **III-** 5 µg/ml LPS, **IV-** 10 µg/ml LPS, **V-** 20 µg/ml LPS, **VI-** 10 µg/ml LPS followed by M-LD, **VII-** M-HD followed by 10 µg/ml LPS, **VIII-** 10 µg/ml LPS followed by M-HD, **IX-** DMSO followed by 10 µg/ml LPS, **X-** 10 µg/ml LPS followed by DMSO. (A combination of groups in Part I and II; see Table I).

#### 3.3.2. *d*-SH-SY5Y Cells

Morphological assessments showed prominent findings *d*-SH-SY5Y cell degeneration especially in the center of large cell aggregates due to MPTP treatment, although cell proliferation and aggregate formation were not affected (Table IV, Figs 7,8).

**Figure 8.**
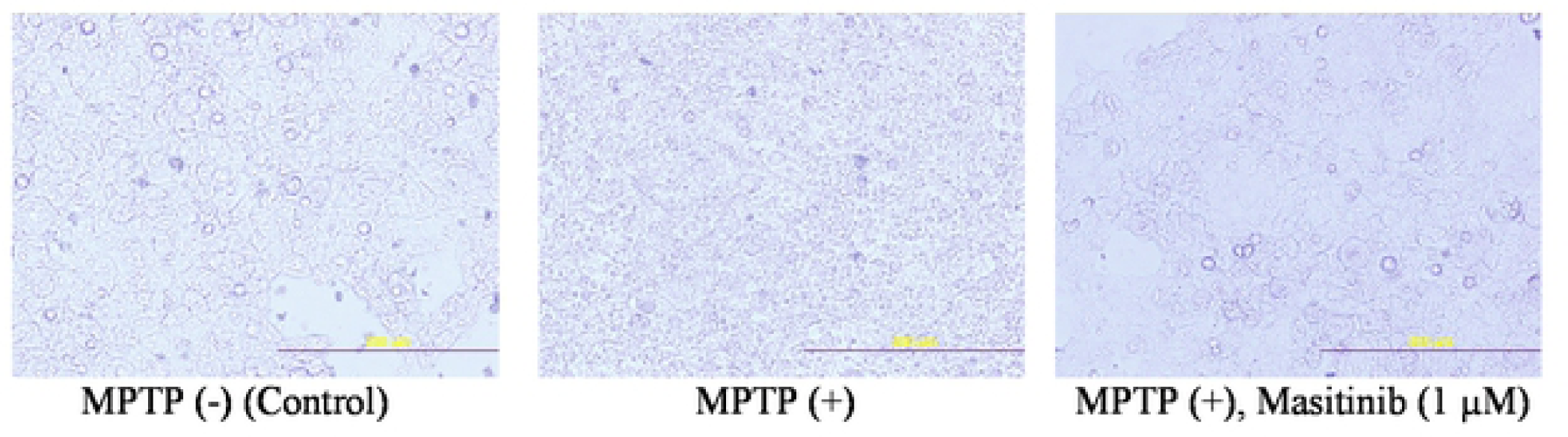
*d*-SH-SY5Y cell culture images showing degenerated cells in MPTP treated group, which was ameliorated with masitinib treatment. (200x), Scale Bar: 200 µm

When compared to control (I), MPTP treated group (II) exhibited increased cell degeneration, whereas masitinib treatment diminished cell degeneration (III).

Activated microglia CM treatment caused cell degeneration (VI-IX), whereas masitinib-treated activated microglia CM caused a lesser percentage of cell degeneration (X, XI).

**Table IV.** Morphological analysis of *d-*SH-SY5Y cells. **I**-MPTP (-) (Control), **II**-MPTP (+), **III**-MPTP (+), Masitinib (1 µM), **IV**-24 hrs non-induced microglia CM, **V**-MPTP (+), 24 hrs non-induced microglia CM, **VI**-MPTP (-), 24 hrs 1 µg/ml LPS-induced microglia CM, **VII**-MPTP (+), 24 hrs 1 µg/ml LPS-induced microglia CM, **VIII**-MPTP (-), 24 hrs 10 µg/ml LPS-induced microglia CM, **IX-**MPTP (+), 24 hrs 10 µg/ml LPS-induced microglia CM, **X-**MPTP (-), 24 hrs 10 µg/ml LPS-induced, masitinib treated microglia CM, **XI-**MPTP (+), 24 hrs 10 µg/ml LPS-induced, masitinib treated microglia CM. (LCAN: Large cell aggregate number, MCAN Medium cell aggregate number, SSAN: scattered small aggregate number, RDILA: The rate of degeneration inside large aggregates, DCS: Degeneration of cells score).

|  | Groups |  |  |  |  |  |  |  |  |  |  |
| --- | --- | --- | --- | --- | --- | --- | --- | --- | --- | --- | --- |
|  | I | II | III | IV | V | VI | VII | VIII | IX | X | XI |
| LCAN | 3 | 4 | 4 | 1 | 1 | 4 | 0 | 3 | 2 | 2 | 3 |
| MCAN | 5 | 2 | 1 | 6 | 0 | 2 | 4 | 10 | 2 | 4 | 8 |
| SSAN | 25 | 40 | 35 | 40 | 45 | 35 | 10 | 55 | 55 | 50 | 50 |
| RDILA (%) | 0 | 40 | 20 | 25 | 50 | 25 | - | 50 | 60 | 40 | 40 |

### 3.4. ELISA and Colorimetric Assay

TGF-β1 levels in microglia CM and *d-*SH-SY5Y cells were measured with ELISA. Moreover NO levels in some microglia CM were measured with a Colorimetric Assay.

#### 3.4.1. TGF-β1 Levels Microglial CM

TGF-β1 levels were found to be silently increased in microglia CM paralel with increasing doses of LPS which was used to induce microglia cells (0-20 µg/ml) (I: 223±32 pg/ml, II: 239±46, III: 299±71, IV: 310±53, V: 377±24 pg/ml). The highest mean level of TGF-β1 was reached in CM of microglia which were activated by 20 µg/ml LPS (Figure 9).

**Figure 9.**
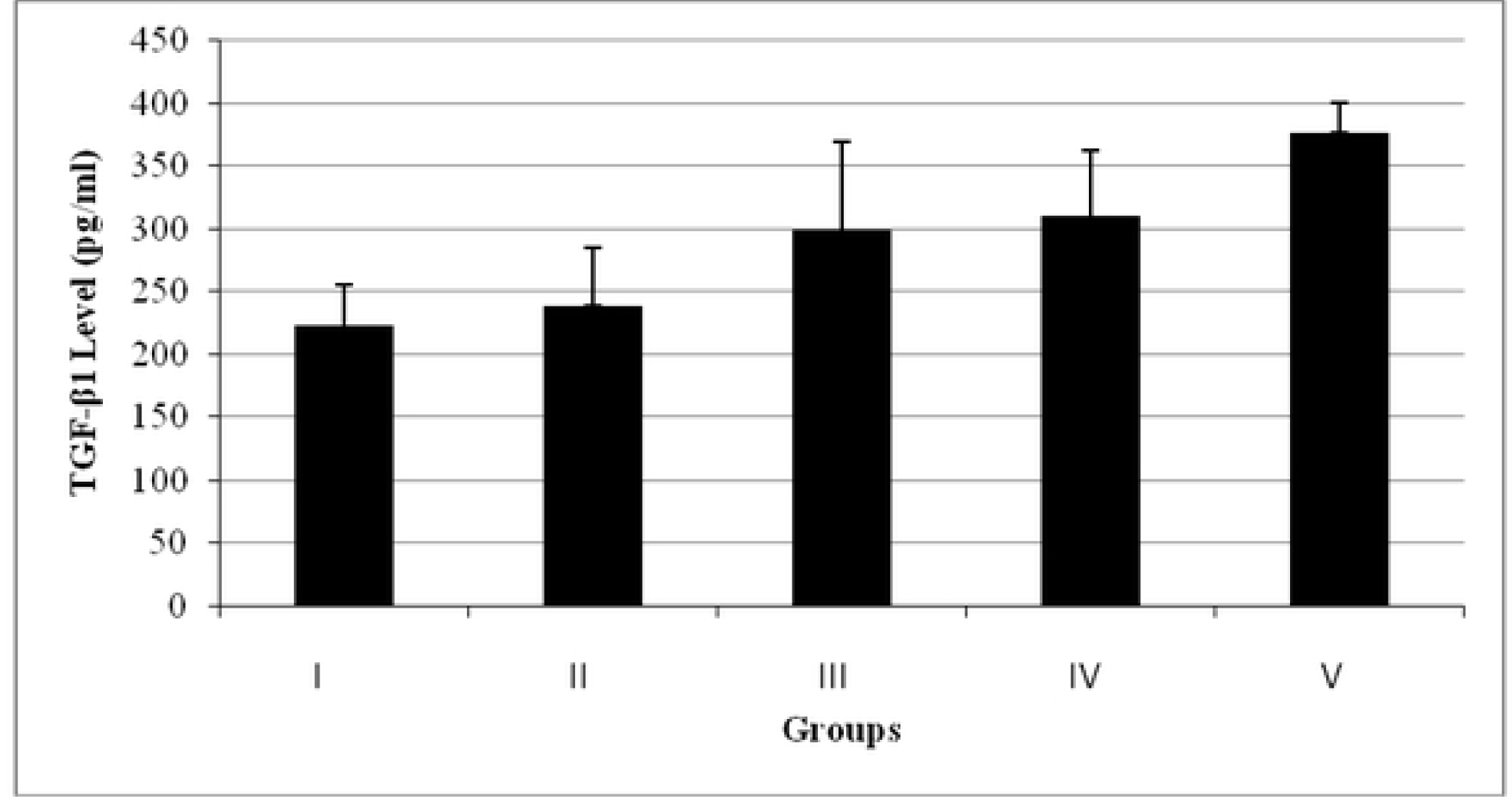
TGF-β1 levels of varying doses of LPS treated microglia CM. **I:** Non-treated, **II:** 1 µg/µl LPS-treated, **III:** 5 µg/µl LPS-treated, **IV:** 10 µg/µl LPS-treated, **V:** 20 µg/µl LPS-treated. Both non-activated and LPS-activated microglia culture CM at 24, 48 and 72 hrs showed an increased level of TGF-β1 in proportion to culture duration (I: 288±35 and 414±42, II: 696±48 and 778±55, III: 895±69 and 934±23, respectively). Note that LPS activated microglia CM showed higher levels of TGF-β1 compared to non-activated ones (Figure 10).

**Figure 10.**
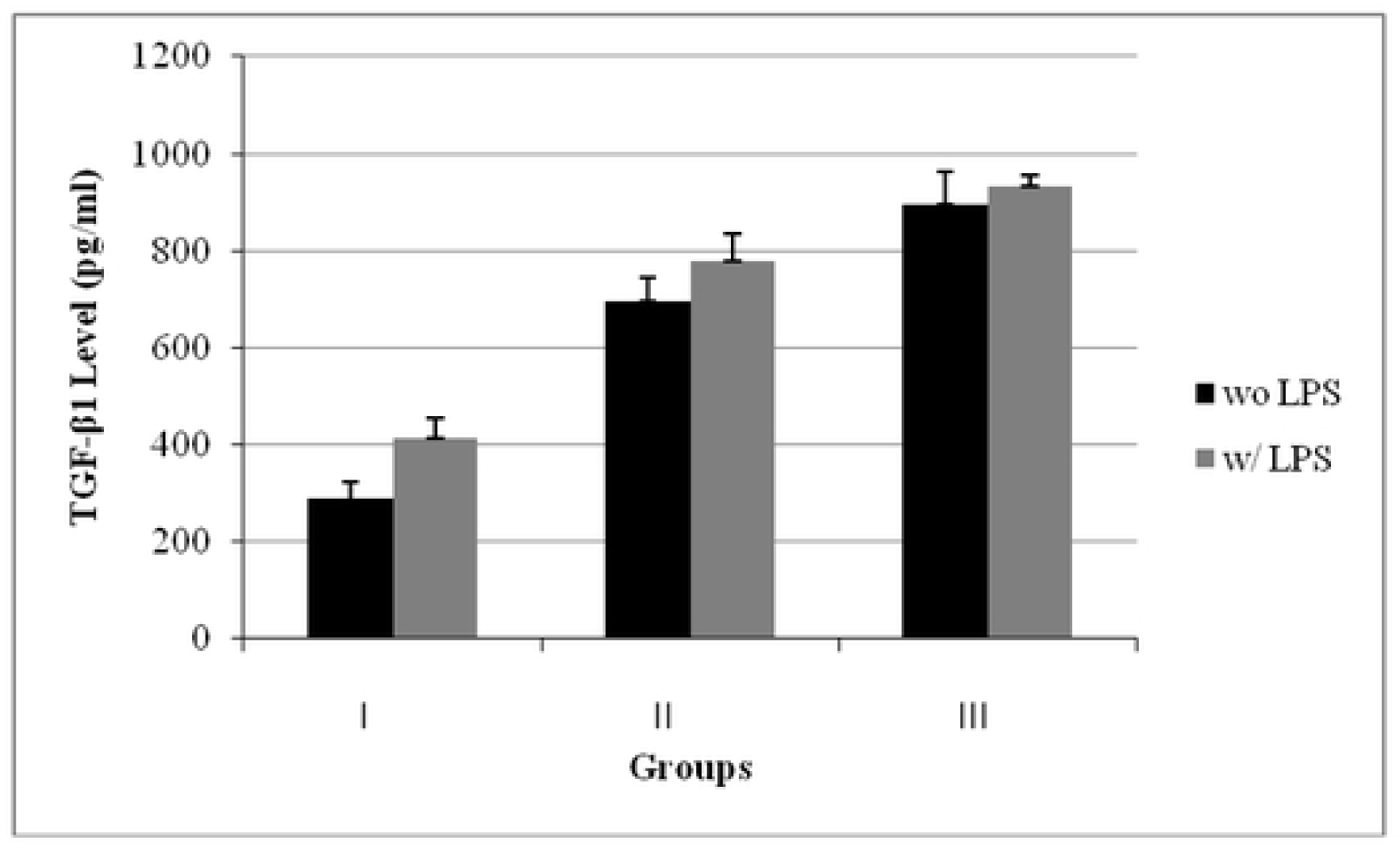
TGF-β1 levels of non-activated or 1 µg/ml LPS-activated microglia CM at different time points. **I:** 24 hrs, **II:** 48 hrs, **III:** 72 hrs When CM of non-induced and 1 µg/ml-LPS-induced microglia which were treated with masitinib were compared in terms of TGF-β1 levels, M-LD provided significant higher levels (III: 694±42 and 500±52 pg/ml) compared to non-treated (I: 288±35 and 414±42 pg/ml) and DMSO treated (II: 243±45 and 274±45 pg/ml) ones (Figure 11).

#### 3.4.2. TGF-β1 Levels in *d-*SH-SY5Y Cell Culture

TGF-β1 levels in CM of non- and MPTP-treated *d-*SH-SY5Y cells (I: 300±48 and 390±36 pg/ml, respectively) were found to be significantly increased when they were treated with M-LD (II: 405±54 and 508±52 pg/ml, respectively). Moreover, when non-activated 24 hrs microglia CM was applied to with or without MPTP *d-*SH-SY5Y cells, TGF-β1 levels were found to be decreased (III: 304±31/423±15 pg/ml, respectively. Furthermore, M-LD treatment before or after this CM application significantly elevated these levels (IV: Early M-LD:730±41/ 663±27 pg/ml; V: Late: 856±15/ 625±15 pg/ml, respectively).

When 24 hrs’ non-activated microglia CM with M-LD was applied to with or without MPTP *d-*SHSY5Y cells, TGF- B1 levels of were found to be significantly increased compared to that of non-masitinib-treated 24 hrs microglia supernatant (VI: 796±22/860±38 pg/ml), although DMSO counterpart of this supernatant caused a prominent decline in TGF-B1 level (VII: 483±10/405±20 pg/ml) (Figure 13).

**Figure 11.**
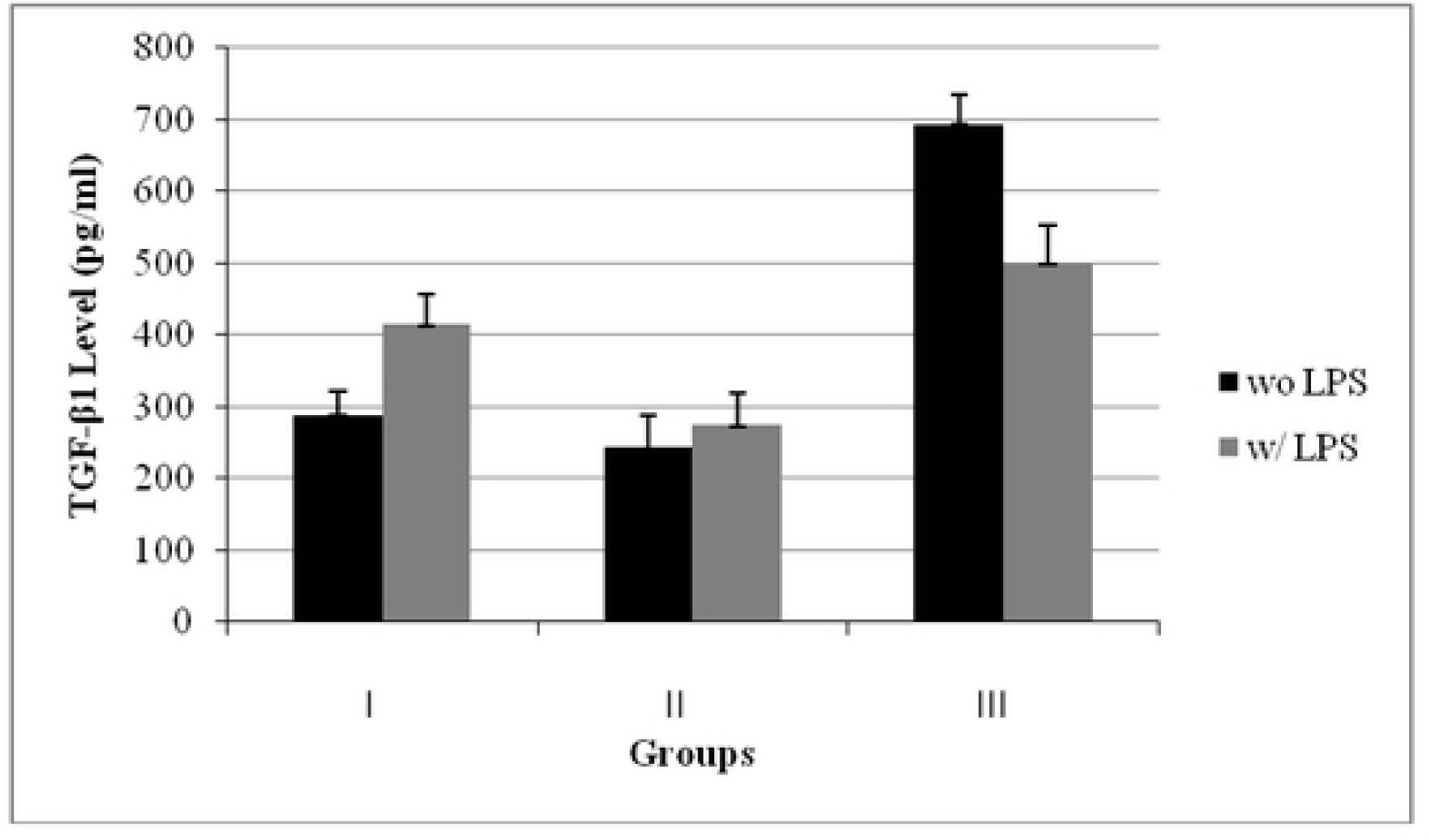
TGF-β1 levels of non-treated, DMSO or masitinib treated microglia (without LPS and 1 µg/ml LPS) CM. **I:** Non-treated, **II:** DMSO treated, **II:** Masitinib (LD) treated When CM of non-induced and 10 µg/ml-LPS-induced microglia which were treated with masitinib/or not were compared in terms of TGF-β1 levels, similiar levels were found in non-treated (I: 223±32 pg/ml), 10 µg/ml-LPS treated (II: 235±53 pg/ml) and DMSO (masitinib solvent) treated (V: 243±42 pg/ml) microglia CM. However, M-LD treatment after 10 µg/ml LPS significantly increased TGF-β1 levels (III: 438±42 pg/ml), whereas M-HD showed a reductive effect (IV: 141±53 pg/ml) (Figure 12).

**Figure 12.**
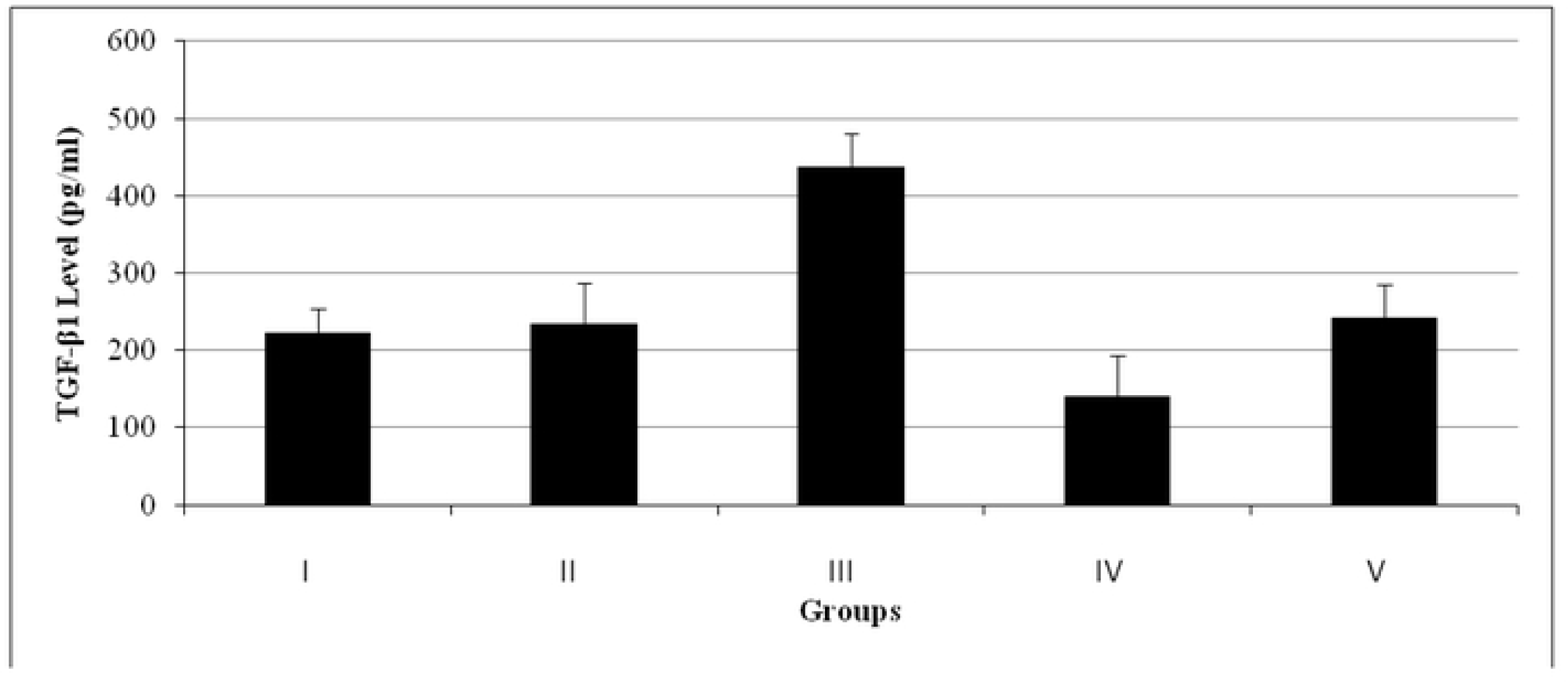
TGF-β1 levels of microglia CM. **I:** Non-treated, **II:** 10 µg/ml LPS, **III:** 10 µg/ml LPS+M-LD, **IV:** 10 µg/ml LPS+M-HD, **V:** 10 µg/ml LPS + DMSO

**Figure 13.**
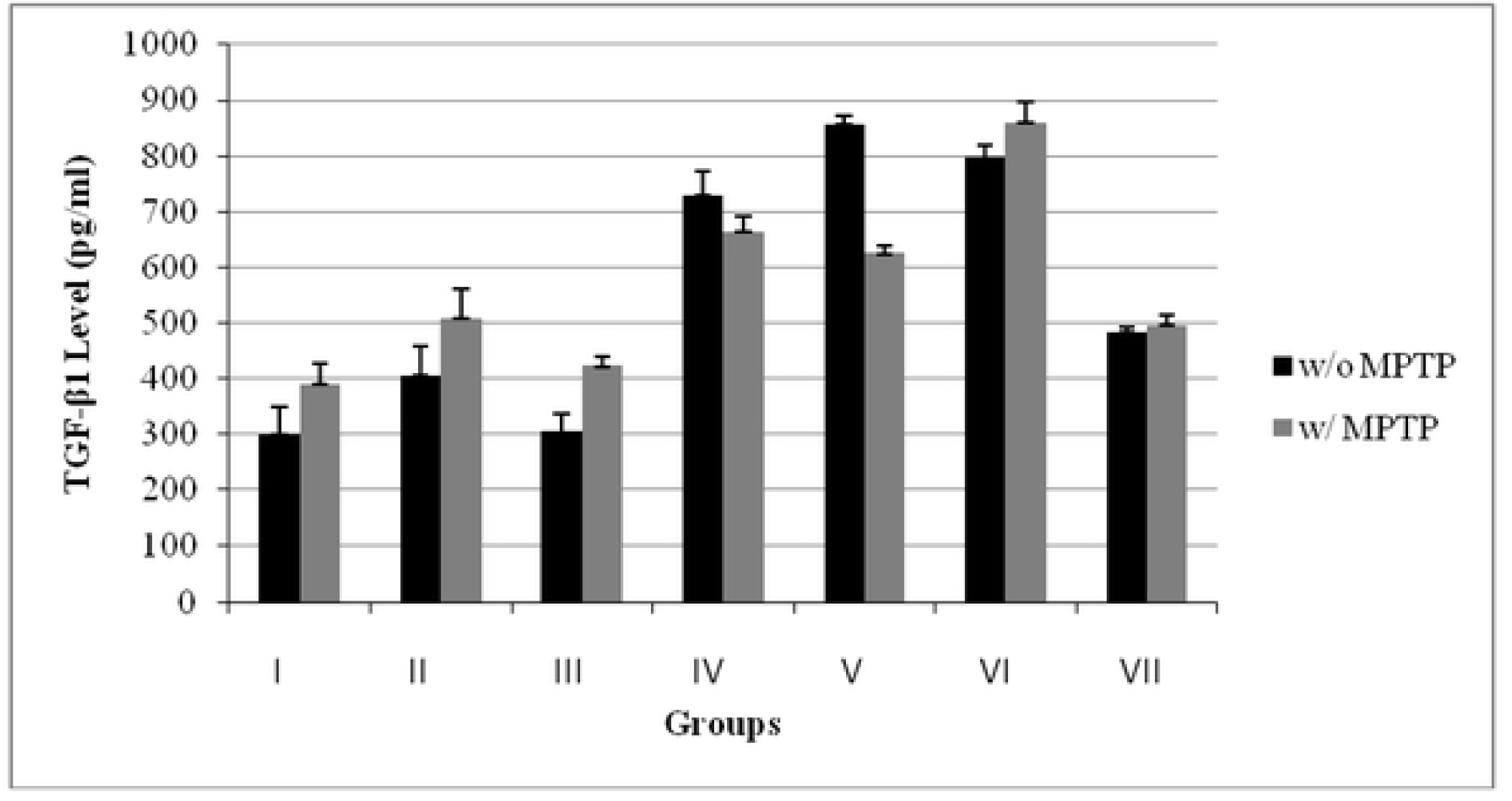
TGF-β1 levels of *d-*SH-SY5Y culture. **I:** Control (with or without MPTP) **II:** M-LD **III:** Non-activated 24 hrs microglia CM **IV:** M-LD (Early) + Non-activated 24 hrs microglia CM **V:** Non-activated 24 hrs microglia CM + M-LD (Late) **VI:** 24 hrs M-LD treated non-activated microglia CM **VII:** 24 hrs DMSO treated non-activated microglia CM

#### 3.4.3. NO Levels in Microglial CM

Higher levels of NO was noticed in CM of microglia which were not activated with LPS (4.3±0.1 µM) compared to that of activated ones with doses of 1, 5 and 10 µg/ml LPS (II: 3.7±0.1, III: 3.6±0.2, IV: 3.6±0.2 µM, respectively).

However, 20 µg/ml LPS- treated (V: 4.8±0.1 µM) group showed higher NO levels compared to the aforementioned groups. Furthermore, M-LD treatment after 10 µg/ml LPS decreased NO levels (VI: 3.3 µM) (Figure 14).

**Figure 14.**
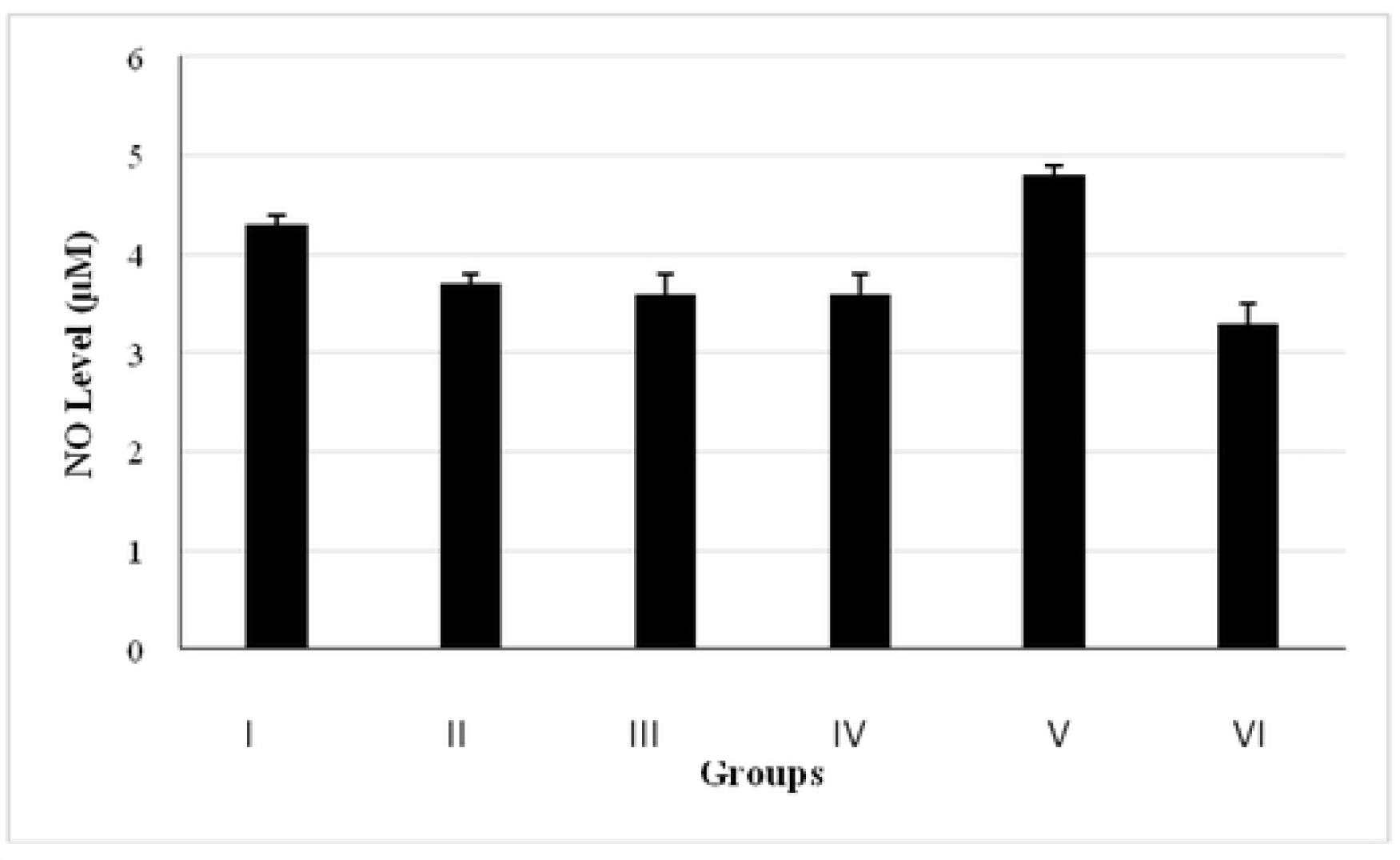
NO levels of CM of microglia treated with M-LD and varying doses of LPS. **I:** Non treated (Control), **II:** 1 µg/µl LPS-treated, **III:** 5 µg/µl LPS-treated, **IV:** 10 µg/µl LPS-treated, **V:** 20 µg/µl LPS-treated **VI:** 10 µg/µl LPS-and M-LD treated microglia CM.

Non-activated or 1 µg/ml LPS-activated groups at 24, 48 and 72 hrs of microglia culture showed different pattens in terms of NO levels. Non-activated microglia CM at 24 hrs of culture (I (wo LPS): 8.5 ±0.2 µM) showed significantly higher NO levels when compared with all the other groups. NO levels at 72 hrs (III: 5.6 ±0.2 µM) in non-activated microglia CM were found to be lower than that of 24 hrs, but higher than that of 48 hrs (4.8±0.2 µM) CM. In activated groups a positive correlation of NO with the duration of culture was observed (Figure 15).

**Figure 15:**
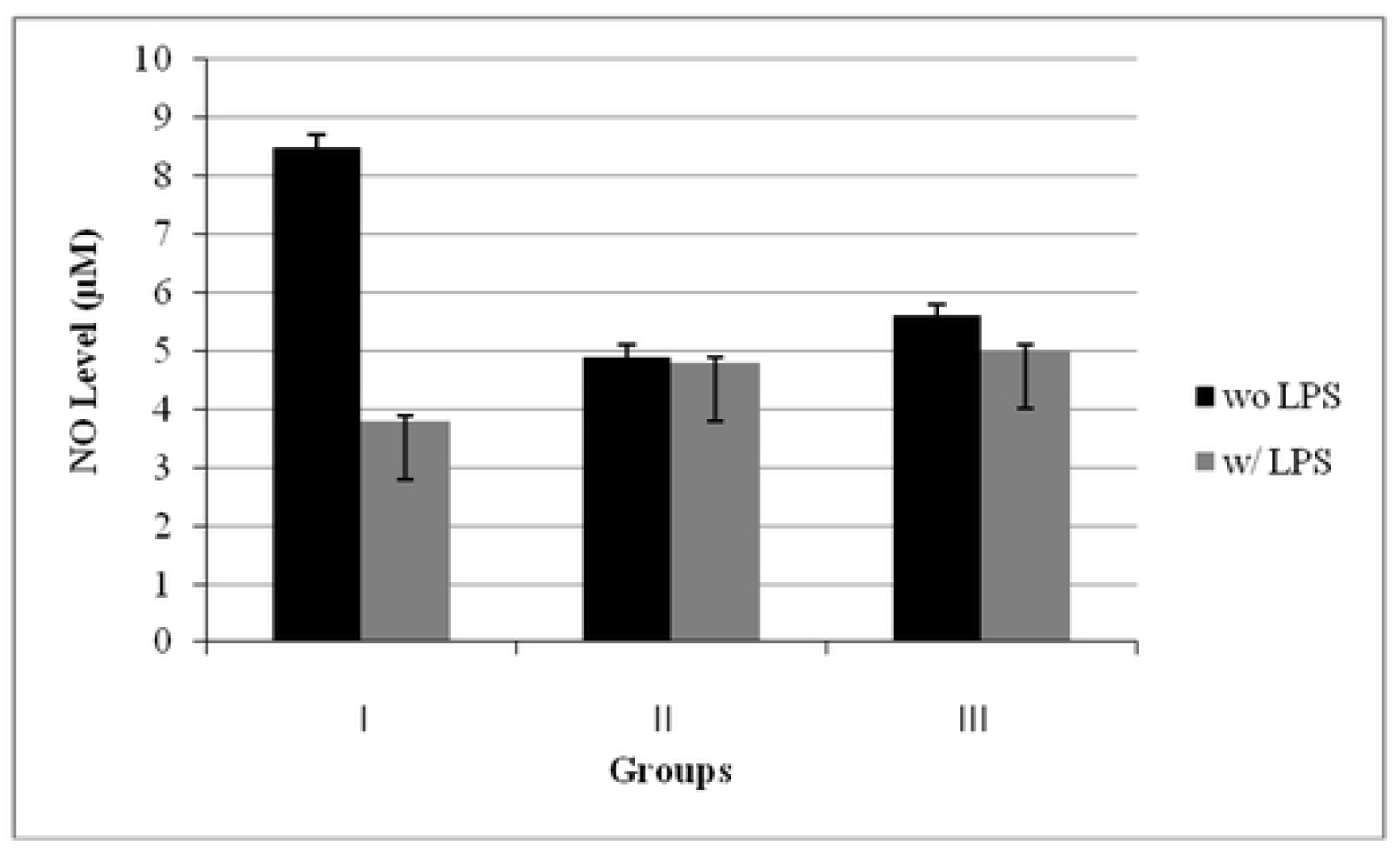
NO levels of non-activated or 1 µg/ml LPS-activated microglia CM at different time points. **I:** 24 hrs, **II:** 48 hrs, **III:** 72 hrs 6-72 hrs culture of non-activated microglia CM showed highest level of NO at 24 hrs (Figure 16).

**Figure 16:**
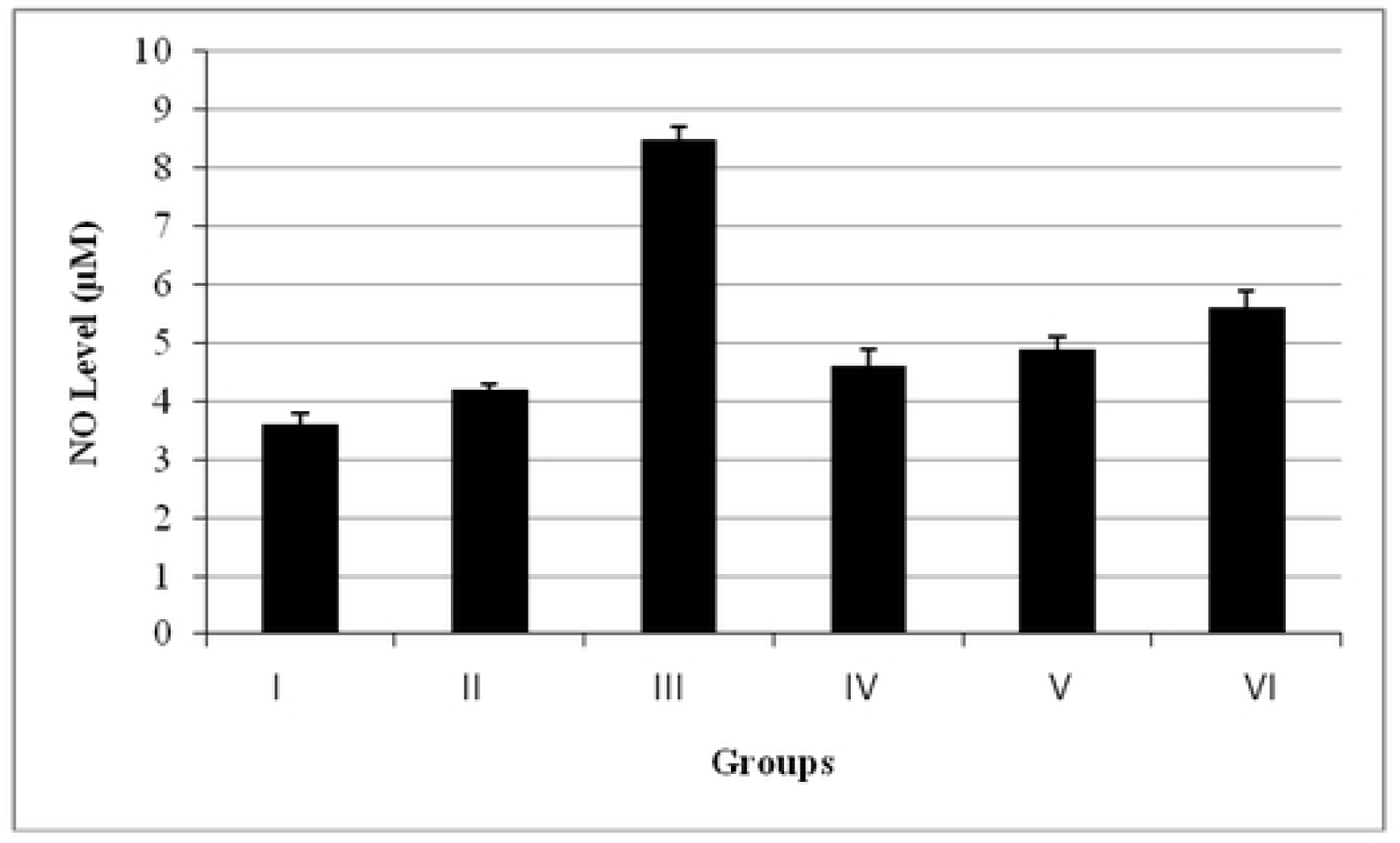
NO levels at 6-72 hrs non-activated microglia CM. **I:** 6 hrs, **II:** 12 hrs, **III:** 24 hrs, **IV:** 36 hrs, **V:** 48 hrs, **VI:** 72 hrs

## Discussion

How neuroinflammation contributes to the progression of neurodegenerative diseases is still unclear as it can be either the cause or the consequence of neuronal cell death. Emerging evidences strengthen the importance of the investigations on neurons to reveal the intracellular mechanisms during neuroinflammation, and their relationship with microglia. Although microglial responses are thought to be of primarily neuroprotective, their roles in tissue injury, neuroinflammation, and neurodegeneration have also been revealed by neuropathological studies. Relevant to their neurodegenerative effect, microglia secrete a wide variety of pro-inflammatory cytokines, toxic enzymes and reactive oxygen / nitrogen species (ROS/RNS) (5,31–33).

In our study we formed an *in vitro* neurodegenerative model by treating *d-*SH-SY5Y cells with MPTP, a neurotoxin that triggers oxidative stress and apoptosis in neurons. MPTP becomes neurotoxic when it is oxidized to an active metabolite, 1-methyl-4-phenylpyridinium (MPP^+^) (34). Once formed, MPP^+^ is taken by dopaminergic cells, where it inhibits NADH-linked mitochondrial respiration, which causes degeneration of dopaminergic neurons (35). MPTP has been shown to cause a mitochondrial damage in SH-SY5Y cells (36). Changes in the cytoskeleton and formation of abnormal neurites have also been observed. Similar changes have been noted in other cell culture systems (37). As survival of neurons are highly dependent on their cellular energy supply and structural integrity, MPTP-induced changes may reflect critical events in neurotoxicity. In this content, our survival analysis demonstrated that MPTP treatment to *d*-SH-SY5Y cells caused a 35% of cell death (Figure 1-5: Control group). This finding was compatible with our morphological assessments, in which a significant degree of cell degeneration was noted especially in the center of large cell aggregates (Figures 7, 8).

Furthermore, we activated microglia HMC3 cells with different doses of LPS, a common inflammatory mediator, and cultured them for different durations to treat their CM on *d-*SH-SY5Y cells. Unraveling the neuroprotective actions of masitinib, we also treated microglia with this agent in different conditions. Levels of TGF-β1, an anti-inflammatory cytokine, and NO, a pro-inflammatory mediator were also determined in microglia CM.

In this context, *d-*SH-SY5Y cells which were treated with CM of microglia -both non-activated and activated-showed decreased rates of cell viability when compared to control. The initial 24 hrs CM of non-activated microglia was found to be the most detrimental medium causing a significant degeneration on with or without MPTP-*d-*SH-SY5Y cells. The second most detrimental CM was from the initial 24 hrs 1 µg/ml-LPS-stimulated microglia. The degeneration increased proportionally with the amount of microglia CM applied.

We also found that non-MPTP treated cells were more susceptible to non-induced microglia CM (Group P1’- I) with a higher rate of cell degeneration compared to that of LPS-induced microglia CM treatment. When the latter CM were applied on *d-*SH-SY5Y cells, the viability of these cells showed a negative correlation with LPS doses that microglia were exposed to (Groups P1’- II-V). In this context, interactions of microglia with themselves and neurons may effectively lead to secretome changes. So, this change may cause different responses to the same stimuli on with or without MPTP- treated cells as in this case.

In contrast to non-treated *d-*SH-SY5Y cells, MPTP-treated cells were found to be more susceptible to LPS-induced microglia, causing increased cell death rates with increasing doses of LPS varying from 1 to 10 µg/ml. However, the viability of *d-*SH-SY5Y cells that exposed to 20 µg/ml LPS group (Group P1-V) demonstrated better survival rates compared with those treated with lower-dose or non-LPS groups (Groups P1’- I-IV). TGF-β1 levels were in parallel with viability results among supernatants of microglia that were activated with increasing doses of LPS (1-10 µg/ml) (Groups P1- II-IV), or non- activated ones (Group P1’- I). However, TGF-β1 levels were significantly higher in CM of microglia which were activated with 20 µg/ml LPS (Group P1-V), than that of aforementioned groups (Table III).

In this context, *in vitro* exposure to LPS and/or interferon gamma (IFNγ) has been associated with morphological alterations of microglia from ramified to amoeboid, an activated or “M1” phenotype which has long been associated with neuroinflammation. LPS mainly initiates the production of pro- inflammatory cytokines such as, TNFα, IL-1β, IL-6, NO, and ROS in both glial cells and neurons (15,38,39). Microglia can switch phenotype when exposed to specific growth factors or cytokines, as well.

Conveniently, in our study, we also found that amoeboid cell count and intensity of CD-11b, a microglial activation marker, showed positive correlations with increasing doses of LPS that microglia were exposed to. Our *in vitro* neurodegenerative model may reflect and mimic the findings of neurodegenerative diseases such that microglia that surround plaques in AD change their morphology from ramified to amoeboid and stain positive for activation markers (40,41). In addition, large numbers of activated microglia can be observed in the CNS and spinal cords of human ALS patients and mouse models (42,43). So, this model may be a good reflection of neurodegenerative diseases and may reflect the *in vivo* environment to some extent making it suitable for drug trials.

Furthermore, we aimed to investigate the protective and therapeutic effects of masitinib on microglial CM-induced degeneration of *d-*SH-SY5Y cells. We have performed masitinib experiments in two different doses of LPS (1 and 10 µg/ml), and mainly two doses of masitinib were administered on microglia; M-LD and M-HD. Both early and late administration of M-LD (Group P2’- III, Group P3’- V and Group P3’- VI) ameliorated the degenerative effects of the related CM in both LPS doses. In contrast to M-LD treatment, when M-HD (Group P3’- VII, VIII, respectively) was administered to 10 µg/ml LPS stimulated microglia, their supernatants did not show better viability rates compared to non-M-HD treated cells, although groups that exposed to CM of 1 µg/ml LPS applied microglia have benefited from M-HD treatment. Conveniently, M-LD treatment to microglia after 10 µg/ml-LPS significantly increased anti-inflammatory TGF-β1 levels in their CM (Group P2-II: 419), although M- HD treatment caused a significant decline in this cytokine level (Group P2-IV: 77).

The treatment of *d*-SH-SY5Y cells with microglial supernatants in which masitinib-solvent, DMSO was administered on activated/non- activated microglia (Groups P3’- IX and X, respectively) resulted in a lesser percentage of cell viability when compared to masitinib-treated ones. We suggest that masitinib ameliorates cell viability by not only reversing the degenerative effects of its solvent, DMSO, but also preventing the degenerative effects of microglial secretions, as well (Figure 3). Proving this, when non-induced microglia which were treated with DMSO were compared with non- treated ones in terms of TGF-β1 levels, it was found that DMSO significantly decreased the levels of this cytokine (325 pg/ml). Despite that, masitinib which applied to non-induced microglia provided significantly higher levels of TGF-β1, suggesting that masitinib reverses adverse effects of DMSO, besides its own enhancer effect on TGF-β1.

Moreover, pro-inflammatory NO levels decrease, whereas TGF-β1 levels increase in microglia CM with increasing doses of LPS. However, 20 µg/ml-LPS treated microglia CM showed high levels of both mediators. Conveniently, this LPS dose was found to be less effective in terms of activating microglial proliferation compared to the effects of 5 or 10 µg/ml LPS treatment, suggesting that LPS may not be effective when applied at very high doses.

Furthermore, we found that the protective effects of masitinib in both with or without MPTP treated cells were significantly better in its 0.5 µM (M-LD) dose of administration, when compared to that of 1 µM (M-HD). While offering a measure of protection, masitinib administration to microglia culture before or after LPS stimulation did not show any significant difference in terms of *d-*SH-SY5Y cell viability (Figure 2).

For both non-activated and 1 µg/ml LPS-activated microglia, the toxicity of supernatants decreased in both 48 and 72 hrs of culture, as proven by cell survival analysis. Improved viability rates of *d-*SH-SY5Y cells in prolonged cultures may indicate that microglial secretions are probably more detrimental at the first 24 hrs of microglial culture (Figure 4). Supporting this, survival rates of *d-*SH-SY5Y cells exposed to wo-LPS /w-LPS groups at 24, 48, and 72 hrs of microglia cultures showed the same pattern with that of TGF-β1 levels. Actually, prolonged culture may lead to up or down-regulation of the various components of the microglia secretome from less than two fold to several thousands-fold by changing the extracellular matrix as well as activating the microglia (6). Moreover, additive and synergistic neurotoxic effects may lead secretome changes as the duration of cell culture prolongs.

When almost all the other groups of microglia supernatants were applied to *d-*SH-SY5Y cells, their survival rates also showed a positive correlation with TGF-β1 levels of the related supernatants (Table III). It suggests that decreased level of anti-inflammatory TGF-β1 may be one of the factors accounting for the decreased viability rates, besides several other cytokines. Conveniently, the expression of TGF-β1 and its receptors were up-regulated in amoeboid microglial cells following hypoxic exposure indicating an autoregulation of microglia in neuropathologies, as reported by Li et al (Li et al., 2008). Additionally, binding of TGF-β1 to its receptors was shown to inhibit free radical production and proliferation of microglia (44–46).

Furthermore, TGF-β superfamily, comprising of TGF-β1-3, has anti-inflammatory action and is generally present in low levels in the brain until there is an inflammation (47). In inflammed CNS, microglial cells are known to secrete TGF-β1 which is involved in regulation of cell growth, differentiation, angiogenesis, and immune function (48,49). Additionally, TGF-β1 was shown to block LPS induced lysosomal acid phosphatase, IL-1β, IL-6 and TNF-α (49), and also induce apoptosis of microglia in a Bcl-2-independent mechanism (50). Thus, there is a subset of evidence supporting the anti-inflammatory function of TGF-β in the CNS and that it plays an important role in neuroprotection (9,15,51)

Finally, to evaluate the direct effects of masitinib treatment on MPTP-related *d-*SH-SY5Y cell degeneration, we also exposed these cells to two doses of masitinib (M-LD, M-HD), and DMSO before (early masitinib) and after (late masitinib) MPTP treatment. Administration of late M-LD (Group P5’- II) significantly ameliorated the viability rates of MPTP treated *d-*SH-SY5Y cells, reaching their viability rate to 100%, as in non-MPTP-treated cells, indicating a fully protective effect (Figure 5).

We found a weak protection of masitinib against MPTP treatment in early administration, suggesting that DMSO content of this drug with high dosage and long duration (an additional 24 hrs) may have harmful effects on cell proliferation. Conveniently, in morphological analysis of early high dosage masitinib treated *d-*SH-SY5Y cells, it was observed that masitinib could prevent degeneration of cells related to MPTP to some extent, but a full recovery was not provided in this dose. We conclude that 0.5 µM dose of masitinib seems to be the most effective dose (Group P5’- II) (Figure 5), showing a protective effect on *d-*SH-SY5Y cells against degenerative effects of neurotoxins.

Our results are supported by studies of Trias et al. (20). They reported that masitinib is capable of controlling microgliosis and the emergence or expansion of aberrant glial cells in the degenerating spinal cord, and has a potent effect in microglial cell cultures, downregulating proliferation, migration, and inflammatory transcriptional profile, and thus controlling neuroinflammation in ALS. They also reported that masitinib significantly prolonged survival when delivered after paralysis onset, making it suitable for treating ALS. Supporting their findings we have found that masitinib shows its effect mainly on activated microglia by reducing their activation, and on degenerated neurons by preventing the neurodegenerative stimulus, or reverse the degeneration before cells progress to necrosis.

In another study, Trias et al showed that mast cells and neutrophils are abundant along the peripheral motor pathway in ALS. These cells appear to be relevant immune cytotoxic effectors in ALS with the potential to be pharmacologically targeted by tyrosine kinase inhibitor drugs. Trias et al. further reported that masitinib ameliorated the peripheral motor axon pathology in SOD1G93A rats, providing another mode of action to explain the promising therapeutic effect of the drug in ALS (52).

Therefore, when considered that masitinib tends to show its neuroprotective effect as the degenerative effects of over-activated microglia or neurodegenerative stimuli are intervened, ***a biomarker*** is a prerequisite for detecting the appropriate application time of treatment. Thus, a biomarker, which can indicate that microglial activation or neurodegeneration has initiated, would be of great interest for researchers to arrange a treatment protocol -with right timing-for masitinib.

In conclusion, here we highlight the neurotoxic potential of resting and activated microglia, and reveal that these cells remarkably contribute to the increased degeneration of neurons. In our study, we also show that masitinib ameliorates viability of with or without MPTP-*d-*SH-SY5Y cells; it not only reverses the degenerative effects of its solvent, DMSO, but prevents the degenerative effects of microglial secretions and MPTP, as well. Therefore, masitinib can be a therapeutic agent in neurodegenerative diseases, as it directly favors neuron survival, and also indirectly protects neurons by reducing the degenerative effects of microglia/other neurotoxins. Masitinib probably provides its final role in neuronal protection by mediating TGF-β1 and NO secretion and by modulating several other mechanisms.

As there are currently no effective treatments for most of the neurodegenerative diseases, our study is valuable in that it indicates the beneficial effects of masitinib on glial secretome, and direct neuroprotective effects. But many other regulatory mediators and direct or indirect cellular interactions remain to be further examined. As we partially predict in vivo behavior of both cell types in our study, we will further do experiments to confirm our primary observations using acutely isolated cells or organoid or other animal systems in which in vivo biology could be better quantified.

## Acknowledgement

This study was supported by the Scientific Research Council of Akdeniz University (Grant number AU, TKA-2018-3933)

